# Efferent feedback enforces bilateral coupling of spontaneous activity in the developing auditory system

**DOI:** 10.1101/2020.08.12.248799

**Authors:** Yixiang Wang, Maya Sanghvi, Alexandra Gribizis, Yueyi Zhang, Lei Song, Barbara Morley, Daniel G. Barson, Joseph Santos-Sacchi, Dhasakumar Navaratnam, Michael Crair

## Abstract

In the developing auditory system, spontaneous activity generated in the cochleae propagates into the central nervous system to promote circuit formation before hearing onset. Effects of the evolving peripheral firing pattern on spontaneous activity in the central auditory system are not well understood. Here, we describe the wide-spread bilateral coupling of spontaneous activity that coincides with the period of transient efferent modulation of inner hair cells from the medial olivochlear (MOC) system. Knocking out the α9/α10 nicotinic acetylcholine receptor, a requisite part of the efferent cholinergic pathway, abolishes these bilateral correlations. Pharmacological and chemogenetic experiments confirm that the MOC system is necessary and sufficient to produce the bilateral coupling. Moreover, auditory sensitivity at hearing onset is reduced in the absence of pre-hearing efferent modulation. Together, our results demonstrate how ascending and descending pathways collectively shape spontaneous activity patterns in the auditory system and reveal the essential role of the MOC efferent system in linking otherwise independent streams of bilateral spontaneous activity during the prehearing period.

## Introduction

Developing sensory systems generate distinct patterns of spontaneous activity to facilitate self-organization and circuit formation before the onset of sensory experience (Blankenship and Feller, 2010, Kirkby et al., 2013, Leighton and Lohmann, 2016, Shrestha et al., 2018). In the cochleae, groups of inner hair cells (IHC) fire bursts of action potentials that propagate to central auditory nuclei via ascending pathways to coordinate activity throughout the auditory system prior to hearing onset (Tritsch et al., 2007, Wang and Bergles, 2015, Wang et al., 2015, Babola et al., 2018). Although central auditory activity may inherit many properties directly from the periphery, it manifests with distinct patterns and novel features (Babola et al., 2018, Sun et al., 2018, Babola et al., 2020). Consequently, understanding the characteristics of spontaneous activity in the central auditory system may provide insight into the maturation of circuit connectivity and the development of higher-order auditory functions.

During the prehearing period, patterns of spontaneous firing of IHCs change across development and vary by tonotopic position in the immature cochlea (Tritsch and Bergles, 2010, Johnson et al., 2011), feeding variable ascending inputs to central circuits. Furthermore, during this time period IHCs’ activity is modulated by transient efferent feedback from the medial-olivocochlear neurons (Glowatzki and Fuchs, 2000, Simmons, 2002, Katz et al., 2004, Goutman et al., 2005), which depends on a9/a10 nAChRs and coupled short-conductance potassium (SK2) channels (Elgoyhen and Katz, 2012). High variability in the peripheral inputs, combined with high fidelity of local coordinated firing, provides substrates for activity-dependent Hebbian plasticity that can promote maturation of precise spatial maps (Feldman, 2012, Kirkby et al., 2013). However, exactly how ascending and descending pathways interact and shape activity patterns to instruct circuit formation remain largely unexplored.

Recent studies have shown coordinated spontaneous activity in the inferior colliculi (IC) at P6-P8, utilizing SNAP25-GCaMP6s mice with wide-field epifluorescent microscopy (Babola et al., 2018, Babola et al., 2020). The calcium transients, detected with the GCaMP sensor, manifest as band-shaped bursts that align across the expected future tonotopic axis in the IC. Here, we conducted a systematic study of spatiotemporal and correlational properties of *in vivo* spontaneous activity in the IC over the entire prehearing period, ranging from postnatal day 0 to 13 (P0-P13). Our results revealed a changing profile of *in vivo* spontaneous activity and evolving bilateral connectivity in the auditory system that peaks early in the pre-hearing period and slowly declines prior to hearing onset. Intriguingly, the strength of bilateral coupling matched the time course of a transient cholinergic modulation imposed on inner hair cells by the medial-olivocochlear (MOC) efferent system (Katz et al., 2004, Elgoyhen and Katz, 2012, Kearney et al., 2019, Frank and Goodrich, 2018). Mice lacking the a9/a10 nicotinic acetylcholine receptor (nAChR), a necessary constituent of efferent modulation (Elgoyhen et al., 1994, Elgoyhen et al., 2001, Morley et al., 2017), displayed severly disrupted bilateral correlations. Chemogenetic and pharmacological experiments *in vivo*, used to acutely manipulate pre- and post-synaptic components of the efferent circuits, indicate that the MOC system was necessary and sufficient to enforce bilateral coupling. Finally, we observed a significant elevation of auditory thresholds at hearing onset in a9/a10 nAChR knockout mice. Together, our results indicate a profound influence of the MOC system, a descending efferent circuit, in coordinating ascending tonotopic bilateral spontaneous activity throughout the auditory system and promoting normal auditory circuit development.

## RESULTS

### Evolving Spatiotemporal Properties of Spontaneous Activity Across the Pre-hearing Period

We first examined how spatiotemporal properties of *in vivo* spontaneous activity changes from birth (P0) until hearing onset (~P13) using wide-field calcium imaging in the mouse IC at six different ages: P0-P1, P3-P4, P6-P7, P9-P10, P11-P12, and P13 (Figures 1A-B). The IC contains three major divisions: the central nucleus (CNIC), the lateral cortex (LCIC), and the dorsal cortex (DCIC). Only the dorsal parts of the IC, including DCIC and LCIC, are accessible for calcium imaging (Babola et al., 2018, Wong and Borst, 2019). Consistent with previous reports (Babola et al., 2018, Babola et al., 2020), typical spontaneous calcium transients in the dorsal surface of the IC appear as stationary and discrete events that (1) have a relatively homogeneous intensity profile along their major axis, (2) show confined spread along the future tonotopic axis that is approximately perpendicular to the major axis (Figures 1C and Supplementary Figure 1B, major axes labeled as magenta arrows), and (3) appear as a single band in the most medial part or dual bands in more lateral parts of the IC (See also Movie S1). We refer to these calcium transients as “spontaneous bands” or “spontaneous events”. We developed a suite of analysis tools for automatic and quantitative description of the spatiotemporal properties of the spontaneous bands (see Methods). We specifically characterized the activity level and spatial profile of spontaneous bands across different ages (Figure 1D-F). Spontaneous bands were present immediately after birth (P0-P1), though with relatively small peak amplitude, and low event frequency. This is consistent with the immature cochlear machinery at P0-P1 (Tritsch and Bergles, 2010). At P3-P4, bands appeared with larger peak amplitude and higher event frequency but had wider spans along the future tonotopic axis compared to bands observed at later stages (Figure 1F). At P6-P7, event frequency continued to increase, indicating that spontaneous activity reached a more active state, and the highest event frequency was recorded at P11-P12, right before hearing onset. We also observed that inter-peak intervals (IPI) decreased monotonically from P0-P12 (Supplementary Figure 1D), consistent with the monotonic increase of event frequency (Figure 1E). Note that spontaneous events at P11-P12 also manifested with more confined width (Figure 1F) and shorter average duration (Supplementary Figure 1E) compared to earlier ages, suggesting that spontaneous bands were more concentrated both spatially and temporally. Spontaneous activity started to diminish at P13, as peak amplitude and event frequency dropped significantly, signaling the end of internally generated activity and the beginning of external inputs (Figure 1D-E). Taken together, spontaneous activity becomes more frequent and spatially refined from birth until hearing onset *in vivo*.

**Figure 1.**
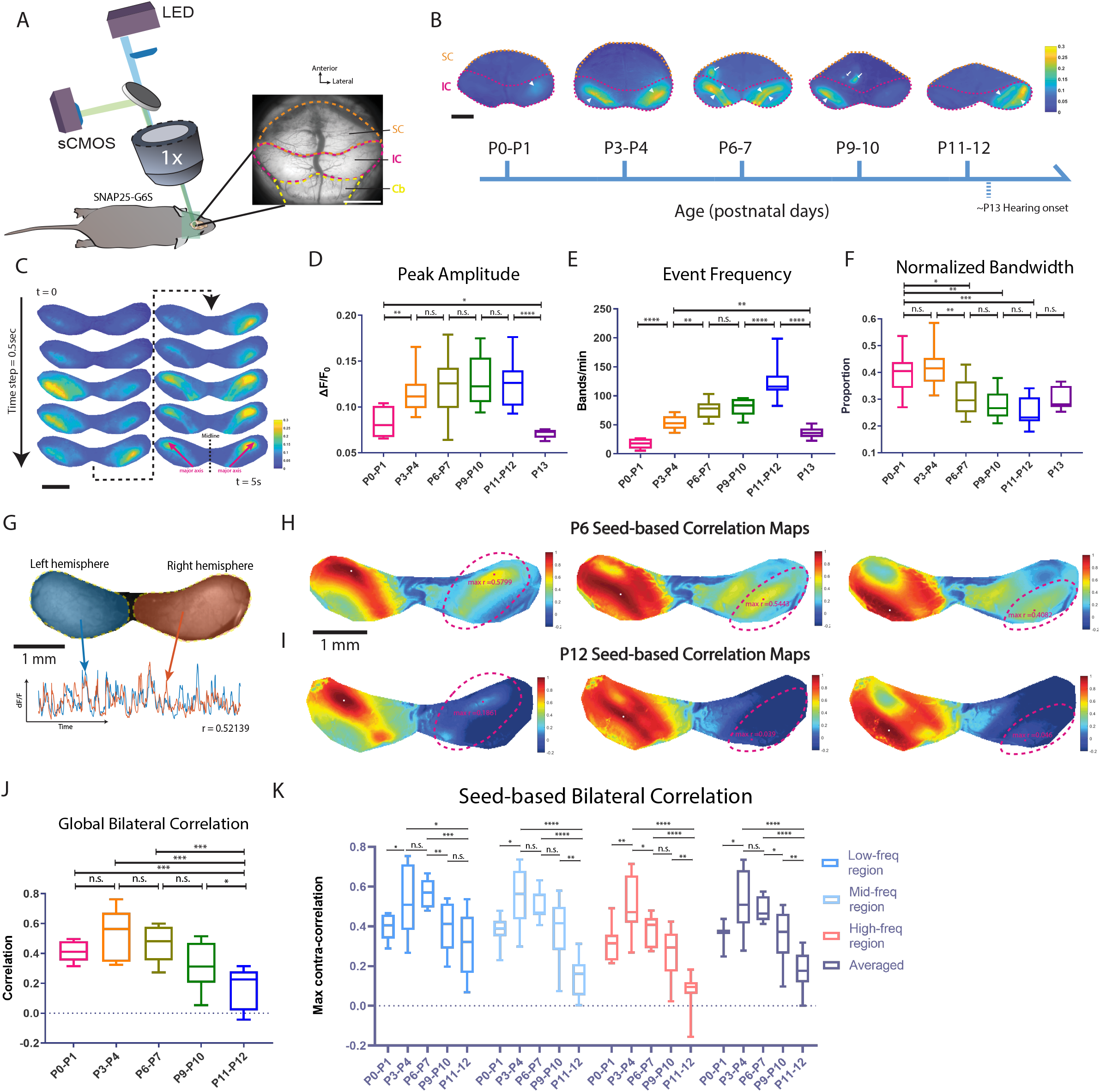
Spatiotemporal and correlational properties of pre-hearing spontaneous activity in the inferior colliculi. (A) Experimental setup of wide-field calcium imaging. Showing a typical field of view of the mouse. Superior colliculi (SC), inferior colliculi (IC), and part of cerebellum (Cb) are delineated in dashed orange, magenta, yellow lines, respectively. All movies are acquired at 10 Hz. (B) Example ΔF/F_0_ images showing spontaneous events at different postnatal ages. Superior colliculi (SC), inferior colliculi (IC) are delineated in dashed orange and magenta lines, respectively. White arrowheads: spontaneous bands in the IC. White arrows: spontaneous (retina) waves in the SC. Colormap: parula (MATLAB). (C) Montages of example ΔF/F_0_ signals appeared as spontaneous bands in the IC. Two magenta arrows represent major axes of the intensity profile in left and right hemispheres. Colormap: parula (MATLAB). (D) Average peak amplitude across age groups. Defined as the mean ΔF/F_0_ amplitude at peaks. (E) Event frequency across age groups. Counted by the number of peaks per minute identified in the line-scan analysis. (F) Average normalized bandwidth across age groups. Defined as the mean spatial half width of fluorescent peaks normalized by the width of the IC. (G) Schematics of global bilateral correlation. Left hemisphere and its corresponding mean fluorescent trace are colored in blue. Right hemisphere and its mean fluorescent trace are colored in orange. Partial correlation between the two traces was computed controlling on the mean activity trace over all pixels outside the IC (r =0.52139). (H) Examples of seed-based correlation maps at P6. Dashed magenta lines delineate symmetrical ROIs in the contralateral hemisphere with respect to the reference seeds. Magenta dots denote maximum correlation in the ROIs (max r). White dots in the left hemispheres indicate seed locations. (I) Examples of seed-based correlation maps at P12. (J) Average global bilateral correlation across age groups (P0-P12). (K) Quantification of seed-based bilateral correlation grouped by future frequency regions. “Low”, “mid”, and “high” correspond to maps where reference seeds locate in the putative low-, mid-, high-frequency regions. “Averaged” correlation is defined as the mean correlation averaged over the three regions. Postnatal ages are shown on the x axis. For all Box plots in this study: hinges: 25 percentile (top), 75 percentile (bottom). Box whiskers (bars): Max value (top), Min value (bottom). The line in the middle of the box is plotted at the median. Significance marks: n.s. p>0.05, * p < 0.05, ** p < 0.01, *** p < 0.001, **** p < 0.0001, two-tailed unpaired t test with Welch’s correction. Number of animals: P0-P1 (N = 8); P3-P4 (N = 11); P6-P7 (N = 10); P9-P10 (N = 9); P11-P12 (N = 11); P13 (N =5). Scale bar indicates 1cm.

We also conducted dimensionality reduction and unsupervised clustering on the calcium data (Supplementary Figure 3). We resolved functional modules that are consistent with known characteristic organization of the IC solely based on spontaneous activity patterns. Specifically, diffusion map results displayed two mirror-symmetrical domains, matching the tonotopic-reversal structure in the dorsal IC (Wong and Borst, 2019). K-means analysis extracted evolving but discrete functional clusters, which are reminiscent of separate fibro-dendritic laminae and discontinuous stepwise response curves in the IC (Jeffery A. Winer, 2005, Malmierca et al., 2008). These results suggest that functional features of the IC are embedded in the intrinsic dynamics of spontaneous activity even before hearing onset.

### Bilateral coupling of spontaneous activity diminishes before hearing onset

Given that central auditory circuits are highly bilateral, we probed correlative features of spontaneous bands in both hemispheres of the IC by examining global bilateral correlations (Figures 1G, 1J) and local (seed-based) correlations (Figures 1H-I, 1K) in the calcium imaging data. After controlling for activity that was external to the selected regions of interests (ROIs; see Methods), background correlations within the IC or between the IC and the visual superior colliculus were reduced, while the salient correlation patterns were preserved (Supplementary Figure 4A, see also STAR Methods). We noticed that global bilateral correlations at P11-12 were significantly lower than at earlier ages (Figure 1J), indicating that average neural activity became less coupled between the two hemispheres at this stage.

The global correlation analysis relies on measurements of activity averaged over the entire IC hemisphere and does not reflect the spatial profile of bilateral correlations or resolve location-specific information. Thus, we also examined spatial correlation patterns in the IC with seed-based correlation maps. We used a high-throughput pipeline to generate numerous maps in parallel, among which we selected respresentative maps for reference seeds located in regions corresponding to future low-, mid-, and high-frequencies (Supplementary Figure 4C). We noticed symmetrical correlation patterns in both hemispheres at P6 (Figure 1H). Specifically, when the representative seed was in the medial IC (putative future low-frequency region), we observed a single correlation band on each side. When the representative seed moved laterally towards the future mid- or high-frequency region of the IC, we observed dual correlation bands that shifted correspondingly to more lateral regions on each side (see Movie S3), reflecting the known reversal of tonotopic organization in each hemisphere (Wong and Borst, 2019). Symmetrical patterns on the two sides of the IC indicate that coupling of bilateral spontaneous activity is itself tonotopic. At P12, however, correlation patterns waned substantially in the contralateral (with respect to the seeds) hemisphere (Figure 1I). We used a novel graphical-user-interface (Supplementary Figure 4B) to quantify maximum correlations within symmetrical regions of the contralateral hemisphere. Summary statistics in Figure 1K showed that seed-based bilateral correlations (referred as SbBC hereafter) increased significantly between P0-P1 and P3-P4 and plateaued through P6-P7. Noticeable drops occurred at later ages, especially at P11-P12, where SbBCs across regions were drastically lower than the level of P3-P7. This increase-peak-decrease trend of bilateral correlations is indicative of developmental changes in functional connectivity between the two sides of the auditory system before hearing onset, which is unexpected given that the anatomy of the mature auditory circuit is highly bilateral. In addition, we noticed a tonotopic difference in bilateral correlations using SbBCs, with low frequency regions exhibiting stronger correlations than those of high frequency regions at P6-7 (Supplementary Figure 4D), suggesting that coupling strengths can vary by tonotopic location.

The spatial profiles of the correlation patterns also provide insight into the local connectivity shaping the spatial properties of the spontaneous activity bands. We observed that the area of the correlations decreased and the eccentricity of the correlation patterns increased over development (Supplementary Figure 4C-E), consistent with decreasing spontaneous activity band widths described in Figure 1F. This suggests that the local connectivity becomes more spatially restricted and laminar-like as animals mature.

In summary, SbBC showed a trend similar to global bilateral correlations, but with more pronounced changes over time (Figures 1J-K). Spontaneous activity in the two hemispheres of the IC has a confined local profile and becomes more uncoupled across different tonotopic regions when maturing towards hearing onset.

### α9/α10 nAChRs are required for bilateral coupling of spontaneous activity

Why does bilateral coupling collapse while activity levels peak at P11-P12? We hypothesize that the medial-olivocochlear (MOC) efferent system might be involved because MOC neurons transiently modulate inner hair cells during the pre-hearing period and are important for normal spontaneous firing patterns (Clause et al., 2014). Previous studies showed that the presence of efferent cholinergic synapses, expression of relevant molecular machinery, and inner hair cell responsiveness to acetylcholine all follow a similar temporal profile to the developmental timeline of bilateral correlations we describe here (Simmons, 2002, Katz et al., 2004, Kearney et al., 2019). Moreover, MOC feedback circuits are known to project to both cochleae. Functionally, ipsilateral sounds can induce both ipsi- and contra-lateral MOC reflexes in mature animals (Liberman et al., 1996, Maison et al., 2003).

MOC efferent axons modulate hair cells through the α9/α10 nicotinic acetylcholine receptors (α9/α10 nAChRs), which are coupled to hyperpolarizing potassium channels/SK2 channels (Elgoyhen and Katz, 2012, see also Figure 2A). In the nervous system, expression of α9/α10 nAChRs is limited to sensory hair cells (Morley et al., 2018). We therefore examined bilateral coupling of spontaneous activity in constitutive α9/α10 knockout mice on the same genetic background (Morley et al., 2017) as the “wild-type” SNAP25-G6s mice. At P6-P7, we saw similar bilateral correlation patterns in the α9/α10 double heterozygous animals (Figure 2B) as in wild-type mice (Figure 1J). However, knocking out α9, α10, or both subunits remarkably impaired correlation patterns in the contralateral hemispheres no matter where the reference seeds were located (Figures 2B-2C, and Movie S4). Note that we grouped the single knockouts (α9/α10 Het and α9 Het/ α10 KO) and the double knockouts (α9 KO/ α10 KO) into the same group (α9/α10 KO) as we observed no significant difference in terms of correlation (Supplementary Figure 6C) or activity levels (Supplementary Figures 6A-B) despite double knockouts showing a trend of more severe bilateral correlation defects (Supplementary Figure 6C). We observed similar event frequencies and a mild increase in peak amplitude in the knockout mice compared to their double-heterozygous littermates (Figures 2D-2E), suggesting that gross activity levels are not compromised in the knockouts. However, we observed a pronounced decrease in bilateral correlations in α9/α10 KOs at P6-P7 (Figure 2F-2G), reminiscent of typical P11-P12 patterns (Figures 1K). Reduced bilateral correlations in the knockouts were also observed at P3-P4 (supplementary Figure 5), indicating an early onset of cholinergic modulation.

**Figure 2.**
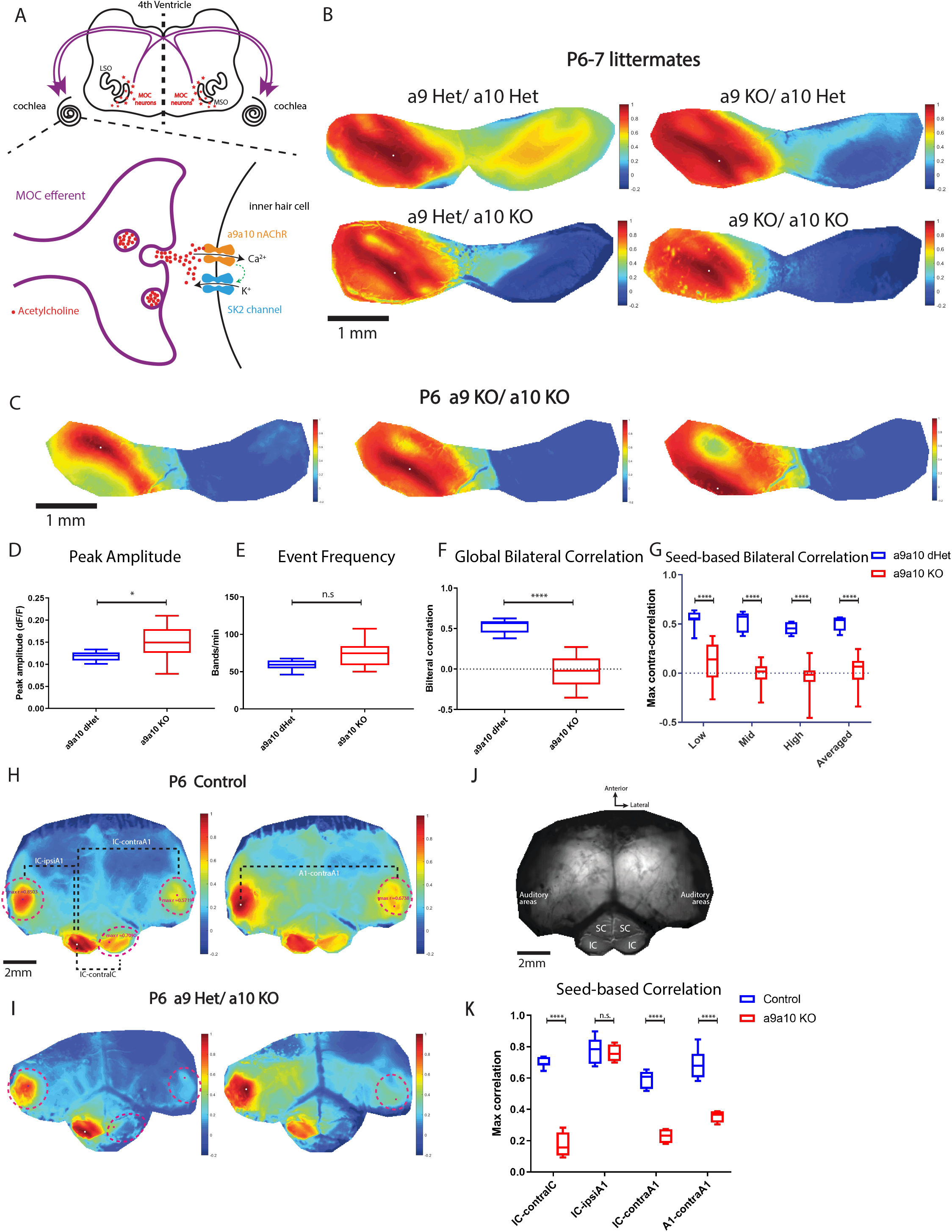
a9a10 nAChR knockouts lack bilateral coupling of spontaneous activity in IC and A1 (A) Top panel: schematics of the medial-olivocochlear (MOC) efferent circuits. LSO: lateral superior olive; MSO: medial superior olive. Purple curved arrows: cochleae receiving bilateral MOC feedback. Bottom panel: simplified schematics of a transient synapse between a MOC efferent axon and an inner hair cell. a9a10 nAChR: alpha9/alpha10 nicotinic acetylcholine receptor. SK2 channel: KCNN2 potassium channel coupled with the a9/a10 nAChR. (B) Example correlation maps showing correlation patterns in the IC among littermates of different genotypes at P6-P7. Het: heterozygous. KO: knockout. Seeds (white dots) are all located in similar regions of left IC. (C) Example IC correlation maps from a P6 a9a10 double knockout animal. Three typical representative seeds (white dots) located in the future low-, mid-, and high-frequency regions. (D) Average peak amplitude. Similar to Figure 1D. (E) Event frequency. Similar to Figure 1E. (F) Global bilateral correlation. Similar to Figure 1J. (G) Seed-based bilateral correlation. Similar to Figure 1K. (H) Example correlation maps with a large field of view including both midbrain and cortex from a P6 SNAP25-G6s animal. Dashed magenta lines: ROIs defined to quantify maximum regional correlations. Left panel: the reference seed is in the left IC (white dot). IC-ipsiA1: between the ipsilateral IC (w.r.t the seed) and the ipsilateral A1. IC-contraA1: between ipsilateral IC and contralateral A1. IC-contralIC: between ipsilateral IC and contralateral IC hemispheres. Right panel: the reference seed is in the left A1 (white dot). A1-contraA1: between the ipsilateral A1 where the seed locates and the contralateral A1. Number of control animals (SNAP25-G6s) = 7. (I) Example correlation maps with a large field of view including both midbrain and cortex from a P6 a9 Het a10 KO animal. Number of a9a10 KO animals = 4. (J) Showing the actual field of view (grayscale mean fluorescent image) over the cortex and the midbrain. SC: superior colliculi. IC: inferior colliculi. Scale bar indicates 2mm. (K) Seed-based correlation. IC-contralIC: between ipsilateral and contralateral IC. IC-ipsiA1: between ipsilateral IC and ipsilateral A1. IC-contraA1: between ipsilateral IC and contralateral A1. A1-contraA1: between ipsilateral A1 and contralateral A1. Box plot symbols similar to (G). (D-G) a9a10 dHet: a9/a10 double heterozygous. a9a10 KO: include single knockout of either a9 or a10 subunit and double knockout of both. Number of animals: a9a10 dHet (N = 7); a9a10 KO (N = 14).

We then used a larger field of view to image bilateral IC and A1 simultaneously in order to examine whether bilateral correlations are similarly altered in developing auditory cortex (Figures 2J). Using seed-based correlation maps, we saw strong correlations between ipsilateral IC/ ipsilateral A1, ipsilateral IC/ contralateral A1, ipsilateral IC/ contralateral IC, and ipsilateral A1/ contralateral A1 in the control (Figure 2H, 2K). In the knockout, however, strong correlations were only present between ipsilateral IC and ipsilateral A1 (Figure 2I, 2K).

In summary, we found that the α9/α10 nAChRs are required for bilateral coupling of spontaneous activity in the IC and auditory cortex. Bilateral correlation patterns were largely absent, as reflected by a significant reduction of correlation values in knockout animals compared to heterozygous littermates. This phenotype was evident from P3-P4 (Supplementary Figure 5) through P6-P7 (Figure 2), when the correlation level peaked in control mice (Figure 1K). These results indicate that the MOC system may play an important role in coupling bilateral spontaneous activity before hearing onset.

### Olivocochlear neurons are necessary and sufficient to mediate bilateral correlations

To investigate whether the loss of bilateral coupling requires acute cholinergic modulation, we directly manipulated cochlear function via topical pharmacological application to the round window (Figure 3A, see STAR Methods). We measured spontaneous activity in control animals (SNAP25-G6s) before and after application of the SK2 channel blocker apamin or the α9/α10 nAChR blocker alpha-Bungarotoxin to both cochleae (see also the schematics in Figure 2A). After application of apamin, correlation patterns in the contralateral IC were largely abolished (Figure 3E), similar to what we observed in the α9/α10 knockout. We compared global and seed-based correlations as well as activity levels before/after pharmacological application (see also Movie S5). We noticed no change in event amplitude or frequency after drug application (Figures 3B-C), while bilateral correlations dropped dramatically in the same animals (Figures 3D and 3F). We next measured bilateral correlations in the IC and A1 in the same animals. Blocking SK2 channel sabolished bilateral correlations while the correlation between ipsilateral IC and A1 was spared (Figure 3G-H), analogous to the phenotypes observed in the α9/α10 knockout (Figures 3H-K). We also observed a similar effect with the α9/α10 nAChR antagonist alpha-Bungarotoxin (Supplementary Figure 7). Applying saline with otherwise identical surgical procedures altered neither correlation profiles nor activity levels (Supplementary Figure 8).

**Figure 3.**
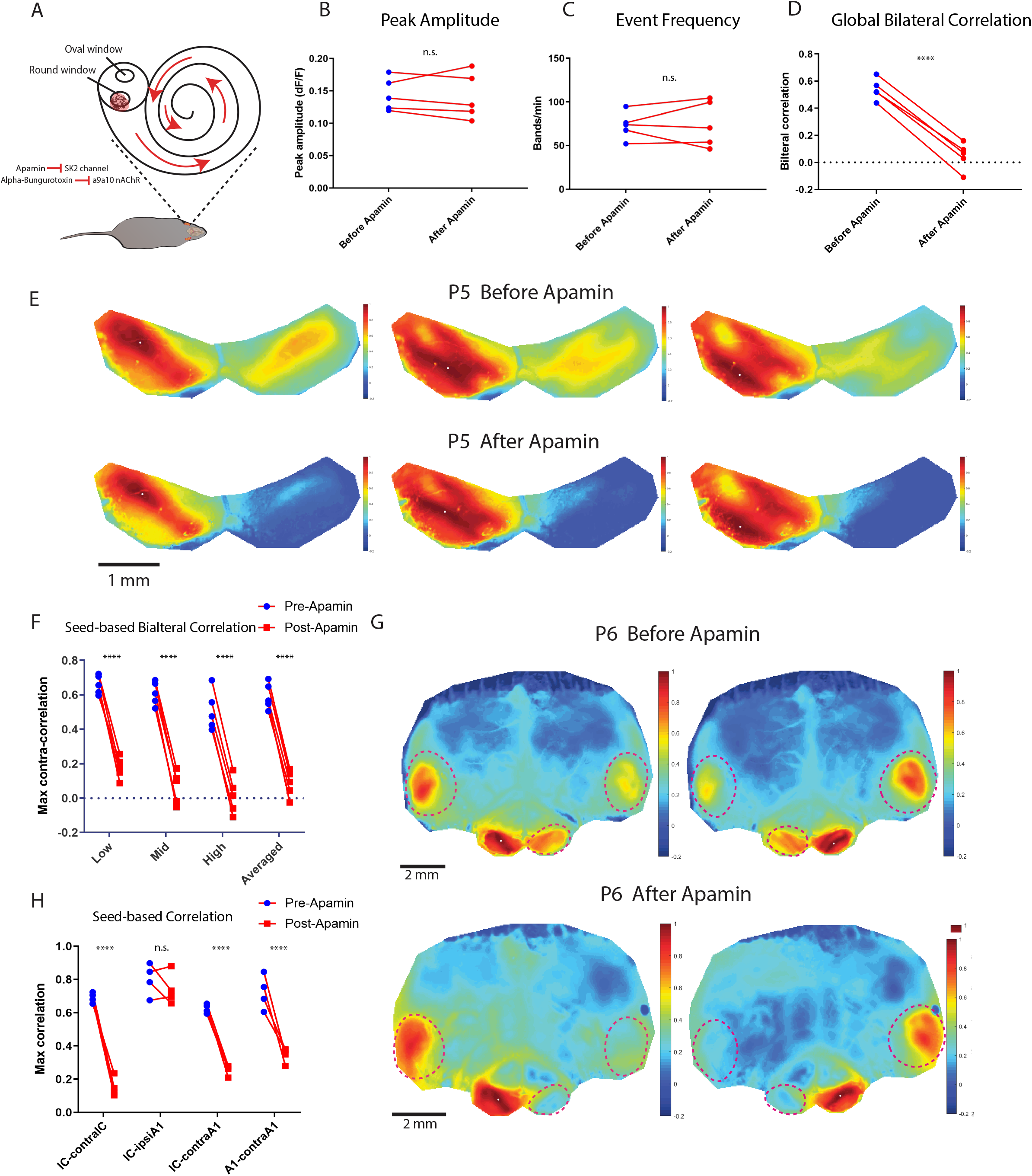
Acute apamin application abolishes bilateral coupling in vivo (A) Schematics showing in vivo pharmacology via round window delivery. (B) Example correlation maps showing correlation patterns in the IC before and after apamin application from the same SNAP25-G6s control animal at P5. Top panels: before apamin. Bottom panels: after apamin. Three typical representative seeds (white dots) located in the future low-, mid-, and high-frequency regions. (C) Average peak amplitude. Similar to Figure 1D. (D) Event frequency. Similar to Figure 1E. (E) Global bilateral correlation. Similar to Figure 1J. (F) Seed-based bilateral correlation. Similar to Figure 1K. (B-F) Number of animals (SNAP25-G6s) = 5. Scale bar denotes 1 mm. (G) Example correlation maps with a large field of view including both midbrain and cortex before and after apamin from the same P6 SNAP25-G6s animal. Top panels: before applying apamin. Bottom panels: after applying apamin. White dot denotes locations of seeds. Dashed magenta lines: ROIs defined to quantify maximum regional correlations. Similar to Figure 2H. (H) Seed-based correlation. IC-contralIC: between ipsilateral and contralateral IC. IC-ipsiA1: between ipsilateral IC and A1. IC-contraA1: between ipsilateral IC and contralateral A1. A1-contraA1: between ipsilateral A1 and contralateral A1. Significance marks: n.s. p>0.05, **** p < 0.0001, two-tailed unpaired t test with Welch’s correction. Number of animals (SNAP25-G6s) = 4.

In summary, our results show that bilateral coupling of spontaneous activity requires cholinergic modulation, presumably from the medial-olivocochlear system, as block of either SK2 channels or α9/α10 nAChRs in the cochlea acutely suppressed bilateral coupling *in vivo*.

To directly probe olivocochlear neurons’ role in coupling bilateral spontaneous activity, we took advantage of cell-type specific chemogenetics (Figures 4A and 4G). We used Designer Receptors Exclusively Activated by Designer Drugs (DREADD)-based chemogenetic tools, which allow suppression or activation of specific neural populations upon application of DREADD agonists (Roth, 2016). Specifically, we injected rAAV-DIO-hM4D-mCherry (inhibitory DREADD) or rAAV-DIO-hM3D-mCherry (excitatory DREADD) retrograde viruses bilaterally in ChAT-CRE;SNAP25-G6s animals via posterior-semicircular-canal (PSCC) injections at P0-P1 (see STAR Methods). Immunohistochemistry confirmed that this approach specifically targeted a subset of ChAT-positive olivocochlear neurons (Supplementary Figure 9). We also verified that the cochlea received inputs from both contralateral and ipsilateral olivocochlear neurons with single-side injections (Supplementary Figure 11).

**Figure 4.**
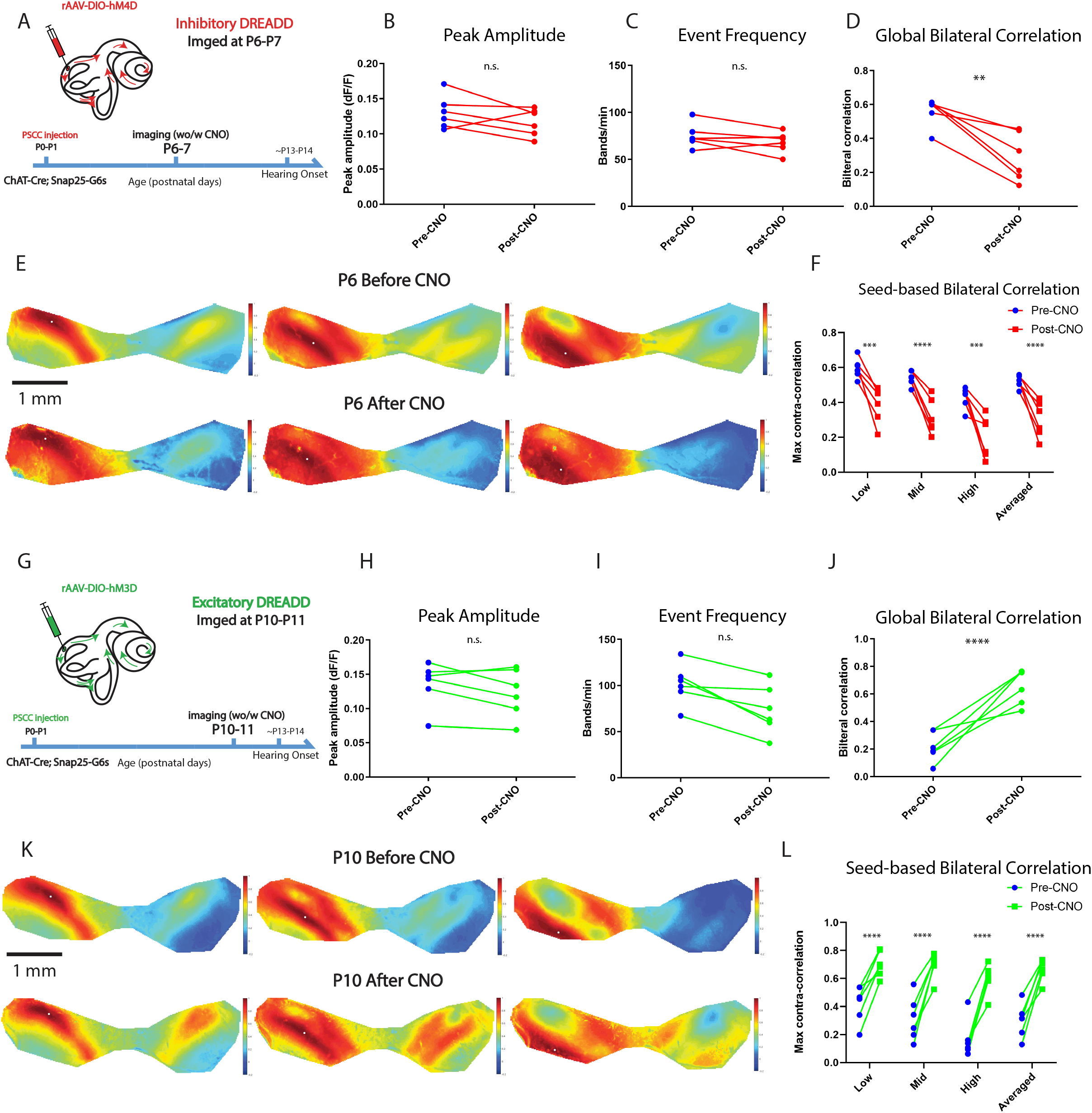
Chemogenetic manipulations can suppress or enhance bilateral coupling in vivo (A) Schematics showing post semicircular canal (PSCC) injection in ChAT-Cre;Snap25-G6s animals and the timeline of inhibitory chemogenetic experiments. (B) Average peak amplitude. Similar to Figure 1D. (C) Event frequency. Similar to Figure 1E. (D) Global bilateral correlation. Similar to Figure 1J. (E) Example correlation maps showing correlation patterns in the IC before and after CNO injection from the same animal at P6. Top panels: before CNO. Bottom panels: after CNO. Three typical representative seeds (white dots) located in the future low-, mid-, and high-frequency regions. (F) Seed-based bilateral correlation. Similar to Figure 1K. (G) Similar to (A) but imaged at P10-P11 for excitatory chemogenetic experiments (H) Similar to (B) for excitatory chemogenetic experiments (I) Similar to (C) for excitatory chemogenetic experiments (J) Similar to (D) for excitatory chemogenetic experiments (K) Similar to (E) but with an example animal at P10 for excitatory chemogenetic experiments (L) Similar to (F) for excitatory chemogenetic experiments Inhibitory DREADD experiments: number of animals = 6; theme color: Red. All imaged at P6-P7. Excitatory DREADD experiments: number of animals = 6; theme color: Green. All imaged at P10-P11.

We first aimed to suppress bilateral coupling with Clozapine N-oxide (CNO), a DREADD agonist. We imaged the animals at P6-P7 when baseline correlation were highest (Figure 1K). Peak amplitude and event frequency (Figures 4B-C) did not significantly change after CNO injection, while global and seed-based bilateral correlations reduced significantly (Figures 4D and 4F). Correlation patterns in the contralateral hemisphere were also impaired (Figure 4E) as in the α9/α10 knockout or pharmacology experiments (Figure 2, 3). These results indicate that olivocochlear neurons are necessary for bilateral coupling *in vivo*.

We then aimed to “boost” the bilateral coupling with CNO. We imaged animals at P10-11 when baseline correlations were low (Figure 1K). Although some MOC-IHC synapses start to degrade at this stage (Elgoyhen and Katz, 2012), MOC mediated modulation can still be observed in IHCs *in vitro* (Kearney et al., 2019). We therefore subjected the olivocochlear neurons to chemogenetic hyper-activation with excitatory DREADDS to enhance efferent modulation. We observed no significant change in activity levels after CNO injection (Figures 4H-I), however both global and seed-based bilateral correlations increased to original peak responses (Figures 4J and 4L). Correlation bands in the contralateral hemisphere were restored after this manipulation, recapitulating the symmetrical patterns observed at P3-P7 (Figure 4K). This result shows that olivocochlear neurons are sufficient to induce bilateral coupling *in vivo*.

In summary, we demonstrated that bilateral coupling can be eliminated or restored by suppressing or activating olivocochlear neurons via chemogenetic manipulations. Combined with the α9/α10 knockout and pharmacological data, our results indicate that the MOC efferent system is both necessary and sufficient to induce bilateral coupling of pre-hearing spontaneous activity.

### Auditory thresholds are elevated in α9/α10 knockout animals at hearing onset

To investigate whether auditory perception was affected in the α9/α10 knockout animals, we first measured auditory thresholds using the auditory brainstem response (ABR). Only ABR waves I-II were visible at P14.0 and were thus used to identify auditory thresholds (Figure 5B, see also STAR Methods). As ABR waves I-II reflect the collective firing of the eighth cranial nerve and cochlear nuclei (CN), responses in higher auditory nuclei downstream to the CN are difficult to detect with the ABR method. ABR thresholds were consistent with previous reports (Song et al., 2006), and there was little difference in responses in α9/α10 knockout and control animals, except at the lowest frequency (Figure 5A). This result indicates that auditory sensitivity up to the level of the CN is relatively normal in the α9/α10 knockout at hearing onset.

**Figure 5.**
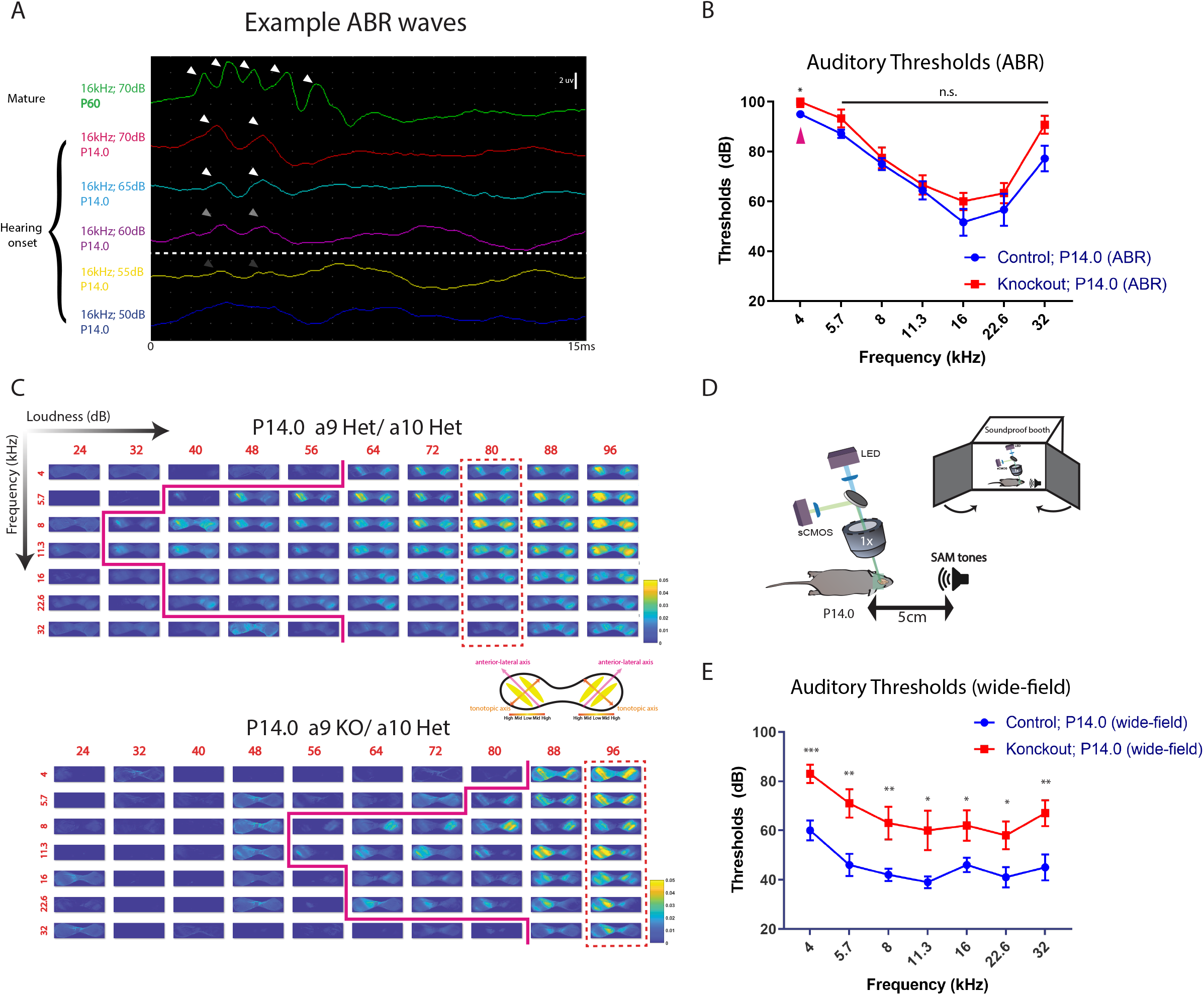
Elevation of auditory thresholds in a9a10 nAChR knockouts at hearing onset (A) Snapshots of example auditory brainstem response (ABR) curves. White arrowheads indicate ABR waves. Scale bar indicates 2uV. White dashed line represents the detection threshold. (B) Auditory thresholds measured by auditory brainstem response (ABR) in control and a9/a10 nAChR knockout animals at P14.0. Number of animals: Control P14.0 = 9 (GCaMP6s negative littermates of aniamls used in (E)). a9a10 knockout P14.0 = 6. Magenta arrowheads indicate that part of the data is not available in the knockout group (two animals did not respond to 4 kHz tones.). n.s. p>0.05, * p < 0.05, two-tailed unpaired t test with Welch’s correction conducted between two conditions at each frequency using available data. (C) Example ΔF/F_0_ response images to different acoustic stimuli. Row direction: loudness. Column direction: frequency. Solid magenta lines denote the auditory thresholds at different frequencies. Dashed red rectangles highlight columns of responses to different frequencies at the same SPL level (tonotopy). Het: heterozygous. KO: knockout. Colormap: parula (MATLAB). Top image array: example from a P14.0 a9a10 double heterozygous control. Bottom image array: example from its P14.0 a9 knockout a10 heterozygous littermate. Schematics in the middle: showing the major axes and tonotopic reversals in the two hemispheres of the IC. (D) Schematics showing the experimental set up of wide-field imaging and acoustic stimulation. The whole apparatus is enclosed in a soundproof booth. SAM: sinusoidal amplitude modulated. (E) Auditory thresholds measured by wide-field imaging. Number of animals: Control = 8 (wide-type SNAP25-G6s = 6; a9a10 double heterozygous SNAP25-G6s = 2). Knockout = 8 (mix of single knockout of either a9 or a10 subunit and double knockout).

To access auditory responses in higher auditory nuclei, we measured auditory thresholds based on activity in the IC using wide-field calcium imaging at P14.0 (Figure 5C, see also STAR Methods). Robust calcium transients in response to acoustic stimuli were present in the IC across the frequency spectrum (Figure 5D and Movie S7). A tonotopic organization of isofrequency bands was observed in both control and knockout animals (columns enclosed in vertical dashed rectangles in Figure 5D), with the medial region of the IC preferentially responding to low frequency, while more lateral regions favored higher frequencies. Typical responses consisted of pairs of bands symmetrically located against the tilted anterior-lateral axes (Movie S7), consistent with the known tonotopic-reversal organization in the IC (Wong and Borst, 2019). Interestingly, we observed a significant elevation on the threshold level of the α9/α10 knockout across the frequency spectrum at P14.0 compared to controls (Figure 5E). Taken together, our results indicate that auditory sensitivity in the α9/α10 knockout is impaired at hearing onset despite minimal changes in the ABR.

## Discussion

*In vitro* studies suggest a system of complex developmental regulation of auditory spontaneous activity in the cochleae (Tritsch and Bergles, 2010). We aimed to examine the developmental profile of pre-hearing spontaneous activity in central auditory circuits *in vivo*. Our experiments revealed the trajectories of key spatiotemporal features of IC spontaneous activity bands throughout the entire pre-hearing period. We observed characteristic organizational features imbedded in the intrinsic dynamics of spontaneous activity in the pre-hearing IC, including discrete functional clusters (Malmierca et al., 2008) and tonotopic-reversals (Wong and Borst, 2019). The large field-of-view afforded by wide-field imaging and the intact *in vivo* preparation permits the direct measurement of functional connectivity across ipsilateral and contralateral hemispheres that are millimeters apart. We observed an unexpected pattern of increase-peak-decrease in bilateral correlations of spontaneous activity between the two IC hemispheres across development. This indicates there are profound changes in bilateral connectivity over the 2-week time window before hearing onset (Figure 1). This developmental timeline suggests that the MOC efferent system might play an important role in coupling bilateral spontaneous activity. Observations in AChR (α9/α10 nAChR) knockout mice, along with pharmacological and chemogenetic experiments (Figures 2-4), confirmed that the MOC system is both necessary and sufficient to enforce wide scale bilateral coupling before hearing onset. These experiments reveal a novel role of MOC efferent modulation on auditory spontaneous activity. Finally, we also observed elevated auditory thresholds in the α9/α10 nAChR knockout mice immediately after hearing onset, based on calcium responses in the IC (Figure 5). Our results, combined with the existing literature (Clause et al., 2014, Clause et al., 2017, Di Guilmi et al., 2019), demonstrate that disruptions of spontaneous activity patterns could undermine different aspects of auditory development.

### Novel aspect of efferent modulation in patterning auditory spontaneous activity

Descending medial-olivocochlear efferent systems have been studied for decades (Warr and Guinan, 1979, Vetter and Mugnaini, 1992, Maison et al., 2003, Ciuman, 2010). In mature animals, outer hair cells (OHC) receive cholinergic feedback from the medial olivocochlear nucleus (MOC). The MOC system suppresses the cochlear amplifier and mediates important functions, such as enhancing signal detection and sound localization in noisy backgrounds, protecting cochlear machinery from loud sounds, mitigating hidden hearing loss, and more (Guinan, 2018). Before hearing onset, however, MOC efferent fibers transiently synapse with inner hair cells (IHC) and modulate IHCs’ spontaneous firing (Glowatzki and Fuchs, 2000, Simmons, 2002, Katz et al., 2004, Goutman et al., 2005). In both cases, this cholinergic modulation depends on α9/α10 nAChRs and coupled short-conductance potassium (SK2) channels (Elgoyhen and Katz, 2012). Previous studies show that animals lacking α9 nAChRs have altered temporal patterns of spontaneous firing (Clause et al., 2014). In this study, we revealed an entirely new aspect of the MOC modulation in patterning spontaneous activity. By taking advantage of the large spatial scale of wide-field calcium imaging, we discovered the MOC’s role in coupling bilateral spontaneous activity throughout the auditory system (Figures 1-4).

These results have several implications. First, bilateral coupling of spontaneous activity may directly instruct bilateral circuit refinement. Central auditory circuits are heavily bilateral at almost all levels. During the prehearing period, peripherally generated spontaneous firing drives patterned activity in the entire auditory system and presumably promotes circuit maturation throughout. However, supporting cells and hair cells in both cochleae cannot communicate directly. Unlike sound waves, which are by nature correlated inputs of the same frequency components, pre-hearing spontaneous activity is independently initiated in the two cochlea. Without the MOC-mediated mechanism that we identified, uncoupled streams of peripheral activity might decrease synchronous activity between presynaptic crossing fibers and postsynaptic auditory neurons, thus undermining bilateral circuit maturation. Moreover, the MOC-mediated bilateral coupling is itself tonotopic, consistent with the known efferent innervation pattern (Frank and Goodrich, 2018). Therefore, the MOC-mediated cochlea coupling can serve as a neural substrate that mimics the bilateral features of real-world stimuli and promotes Hebbian plasticity across two sides of the auditory system. Indeed, Clause et al. (Clause et al., 2014) showed that MNTB-LSO (medial nucleus of the trapezoid body to lateral superior olive) connectivity, where activity is integrated from both cochlea, is impaired in α9 nAChR knockouts, but not in the CN-MNTB calyces, where ipsilateral activity dominates. Behaviorally, these bilateral circuits are the neural basis for critical auditory functions such as sound localization (Grothe et al., 2010). α9 nAChR knockout animals also displayed deficits in frequency processing and sound localization at hearing onset (Clause et al., 2017). Note that previous studies attribute these phenotypes solely to altered temporal patterns of spontaneous activity. We suggest that, though precise temporal patterns may play an important role at the synaptic level, system-wide bilateral coupling informs auditory circuit refinement on a macroscopic scale. Together, we propose that spatiotemporal patterns of spontaneous activity can instruct auditory system maturation by recapitulating natural features of sound. Interestingly, synchronous retinal spontaneous activity (“retinal waves”) were also observed in the binocular zone of the visual midbrain (superior colliculi), possibly mediated by retinopetal or retino-retinal circuits (Muller and Hollander, 1988, Gastinger et al., 2006, Ackman et al., 2012). From an evolutionary perspective, mammalian neural systems might engage in sophisticated regulation of spontaneous activity patterns to prime sensory circuits for future external inputs.

A second implication of these results stems from the developmental timeline of bilateral coupling, which might reflect changes of efferent modulation strength *in vivo*. Specifically, we observed that bilateral correlations peaked at P3-P7 and started to decrease at P9-P10. Bilateral spontaneous activity became even less coupled as correlations approached zero at P11-P12 (Figures 1K). P10-P11 is considered a transition age where MOC-OHC synapses form and MOC-IHC synapses start to decompose (Simmons, 2002, Elgoyhen and Katz, 2012). Indeed, MOC mediated IPSCs can be recorded in both IHCs and OHCs at P11 (Ballestero et al., 2011, Kearney et al., 2019, Vattino et al., 2020). Consistently, bilateral coupling can be rescued with chemogenetic excitation at P11 (Figure 4 J-L), suggesting the existence of MOC-IHC synapses in vivo. On the other hand, acetylcholine release from MOC fibers is negatively regulated by BK potassium channels after P9 and differentially regulated by different types of voltage-gated calcium channels before and after P9 (Kearney et al., 2019). We observed a similar transition stage of bilateral correlation at P9-P10, consistent with the above-mentioned regulatory changes. This suggests that MOC-IHC synapses at this late stage (~P11), though still functional, are less effective in modulating spontaneous activity *in vivo*.

Finally, MOC modulation might differ by tonotopic location. We utilized seed-based correlation maps to visualize bilateral correlation patterns, which also allowed us to differentiate correlation strengths at different future tonotopic locations. Although bilateral correlations showed similar temporal trends over development (Figure 1K), we found that future low-frequency regions were generally more bilaterally coupled than future high-frequency regions starting at P6-P7 (Supplementary Figure 4F). This implies that MOC modulation might be stronger at the apical turn (low-frequency) than the basal turn (high-frequency region) in the cochleae. Indeed, *in vitro* electrophysiology data suggests that spontaneous firing of IHCs at the apical turn are more strongly modulated, presumably by the efferent system, than those at the basal turn (Johnson et al., 2011).

### Reduced auditory sensitivity in the α9/α10 nAChR knockout at hearing onset

In mature animals, MOC efferent fibers inhibit OHCs and suppress cochlear amplification (Ryugo et al., 2011). Here we reported elevated auditory thresholds in the α9/α10 knockout at hearing onset across the frequency spectrum (Figure 5). Although MOC-OHC synapses are already functional at this age (Vattino et al., 2020), the deficit is probably not caused by loss of acute MOC inhibition on the cochlear amplifier. In fact, eliminating the efferent modulation does not change auditory sensitivity as knocking out the α9 nAChR or the coupled SK2 channel does not alter ABR thresholds in mature animals (Ryugo et al., 2011, Morley et al., 2017), while strengthening efferent inhibitions leads to threshold elevation (Taranda et al., 2009). On the other hand, animals should have gained very limited auditory experience at this early age, as auditory thresholds are extremely high before P14.0 (Song et al., 2006) (see also Supplementary Figure 10E). Taken together, our results suggest that threshold elevation in the α9 nAChR knockouts at hearing onset is primarily a developmental ramification. Note that this observation is based on central responses from the IC using wide-field calcium imaging, which are difficult to detect with the ABR method (Figure 5C-E). Thresholds determined with the only visible ABR waves I-II are largely normal in the α9 nAChR knockouts compared to controls (Figure 5A-B), consistent with data from adults (Morley et al., 2017). Our results suggest that central circuits downstream to the CN, rather than the peripheral machinery, are undermined in the knockout, which are in line with previous reports of impaired tonotopic refinement and synaptic functions in the MNTB with either the α9 nAChR knockout (Clause et al., 2014) or the enhanced α9 nAChR knock-in (Di Guilmi et al., 2019). However, more in-depth investigation on the intactness of the sensory periphery is needed for a more definitive conclusion. Nevertheless, accumulated evidence stresses the important role of precise spontaneous activity patterns in facilitating auditory circuit maturation and the development of auditory function.

We observed similar ABR thresholds in control animals as previously reported (Song et al., 2006), but lower thresholds at most frequencies measured with calcium imaging (Supplementary Figure 10E). There are significant differences between the experimental set-ups of the two modalities (ABR and wide-field imaging). Wide-field imaging captures slow calcium activity on the scale of 50-100ms, limited by the dynamics of GCaMP sensors, while the ABR captures fast electrophysiological dynamics on the scale of <1ms. Accordingly, acoustic stimuli to induce calcium responses were usually much longer than those used in the ABR (Zheng et al., 1999, Song et al., 2006, Issa et al., 2014, Romero et al., 2020). In our experiments, animals were fully awake during wide-field imaging of acoustic stimuli, sinusoidal-amplitude modulated tones of 500ms duration were presented, and average calcium responses were acquired over 6-10 repeated sessions; for the ABR, animals were anesthetized, pure tone pips of 3ms duration were presented, and average ABR waves were acquired over 400 repeats. Moreover, although the acoustic systems for the ABR and the calcium imaging were calibrated to have similar response curves, the exact apparatuses and background noise levels were still different (see STAR Methods). Therefore, auditory thresholds measured with the ABR cannot directly translate to those measured with wide-field calcium imaging. However, our results did demonstrate that, (1) central auditory responses in the IC can be detected by calcium imaging while only ABR waves I-II were consistently visible at P14.0 (Figure 5B and 5D), and (2) wide-field imaging could potentially capture lower auditory thresholds at hearing onset based on responses in higher auditory nuclei than the levels previously considered in the field. In fact, decades of psychoacoustic experiments on humans and animals indicate that behavioral thresholds can be ~20-30dB lowers than those measured with the ABR (Ehret, 1977, R.Fay, 1988, Pinto and Matas, 2007, Crowell et al., 2016). Therefore, it is possible that central auditory circuits can pick up signals from just a few peripheral neurons through a cascade of amplification and generate activity detectable with calcium imaging, while remaining elusive to the ABR. Our results suggest that wide-field calcium imaging is a complementary technique for measuring auditory responses, especially in higher order auditory nuclei.

### Potential circuit basis underlying bilateral coupling

Unilateral MOC neurons receive inputs from and provide feedback to both cochleae (Maison et al., 2003, Ryugo et al., 2011). Functionally, presentation of ipsilateral sounds can induce ipsilateral and contralateral MOC reflexes in mature animals (Guinan, 2018, Lopez-Poveda, 2018). We showed that single-side injections of retrograde virus at P0-P1 can target olivocochlear neurons on both sides, confirming the existence of bilateral efferent circuits in neonates (Supplementary Figure 11). The distribution of labeled neurons resembled what had been described as the “DPO (dorsal preolivary)” and/or “shell” neurons (Vetter and Mugnaini, 1992, Brown and Levine, 2008). An important question is: What drives these MOC neurons during the prehearing period? We propose two models here (Figure 6): Frist, MOC neurons may be driven by the cochleae. This is the case in mature animals, where dominant inputs to the MOC neurons ultimately come from hair cells (Brown et al., 2003, Maison et al., 2003, Froud et al., 2015, Maison et al., 2016, Guinan, 2018). Second, there may be other common drives. The MOC neurons are innervated by different types of dendrites, suggesting heterogeneous inputs (Benson and Brown, 2006). A recent study shows that MOC neurons receive inhibition directly from the MNTB (Torres Cadenas et al., 2020). Therefore, it is possible that other auditory nuclei may coordinate MOC neurons in both hemispheres. Note that these two models are not mutually exclusive and could work in synergy.

**Figure 6.**
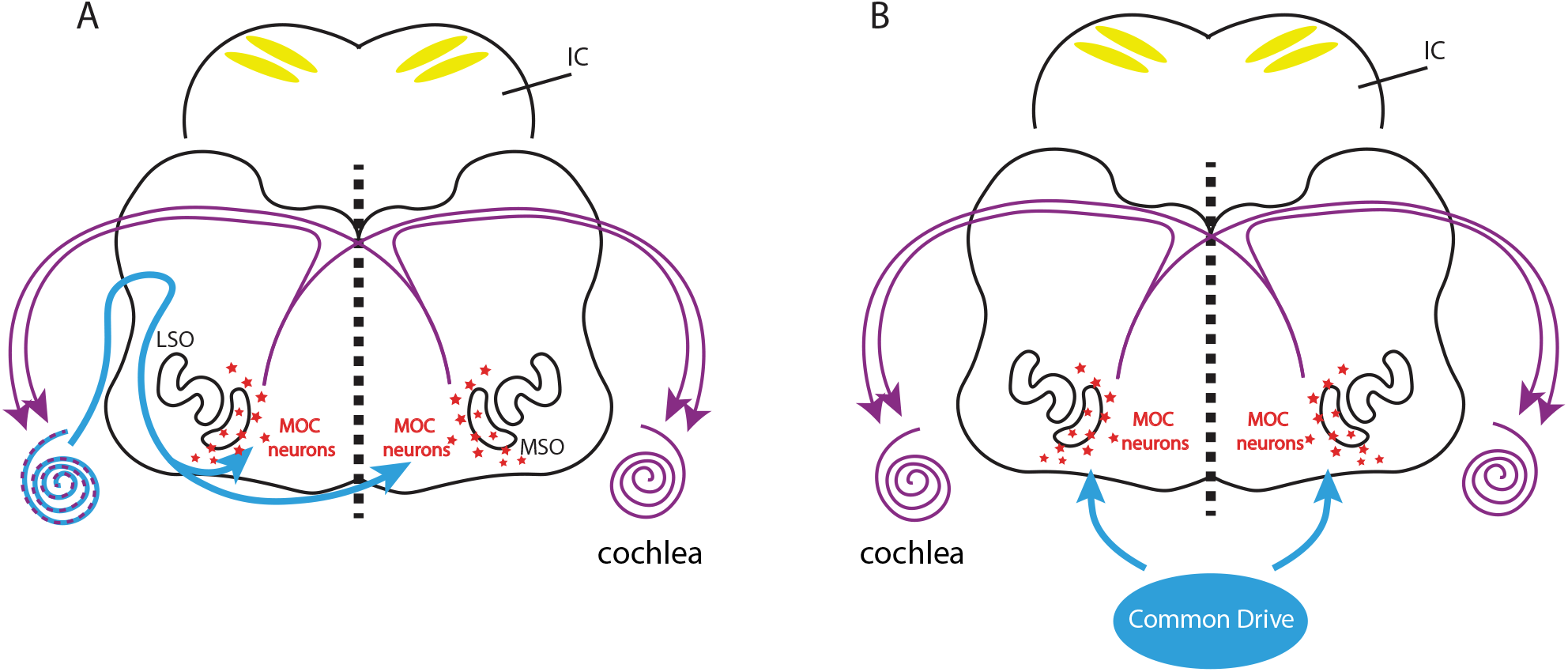
Models of MOC-mediated bilateral coupling of spontaneous activity (A) Model of the medial-olivocochlear (MOC) efferent circuits mediating bilateral coupling of spontaneous activity. Showing the brainstem section where MOC neurons reside. LSO: lateral superior olive; MSO: medial superior olive. Purple curved arrows: cochleae receiving bilateral MOC modulation. IC is shown in the background with bilateral calcium bands presented. Driving inputs from cochleae to the MOC labeled as blue curved arrows. (B) Similar model as in (A). Driving inputs from other common sources to the MOC labeled in blue.

### Auditory spontaneous activity develops in a continuous fashion *in vivo*

Pre-hearing auditory spontaneous activity exhibits different firing properties at different ages *in vitro* (Jones et al., 2007, Tritsch and Bergles, 2010, Yin et al., 2018). In comparison to the step-wise development of the visual system (Ackman et al., 2012, Arroyo and Feller, 2016, Gribizis et al., 2019, Ford et al., 2012), it is intriguing to consider whether auditory spontaneous activity develops in a similar stepwise or continuous/smooth fashion. In the mouse cochlea, supporting cells display increasing firing amplitude, intracellular calcium transient level, and mechanical crenation magnitude from birth until ~P10. Inner hair cells, driven by adjacent supporting cells, show a similar monotonic increase of spontaneous activity levels over the same period. Spontaneous firing in both cell types drops significantly at P13 (Tritsch and Bergles, 2010). Comparable trends are observed in mouse cochlear nucleus (CN) and cat auditory nerves (Jones et al., 2007, Yin et al., 2018). Here, we report similar developmental changes regarding spontaneous bands in the IC, as *in vivo* activity levels increased from P0-P12 and then decreased at P13 (Figure 1D-E). On the other hand, IHCs stay responsive to ATP during the pre-hearing period, suggesting that auditory spontaneous activity might be initiated by the same purinergic machinery before hearing onset (Tritsch and Bergles, 2010). We also showed that spontaneous events in the IC generally displayed band-shape intensity profiles with relatively smooth spatiotemporal variations, rather than crisp transitions across ages (Figure 1 and Movie S1). Taken together, *in vivo* and *in vitro* data suggests a continuous progression of the spatiotemporal features of auditory spontaneous activity during the prehearing period, in contrast to the mechanistically distinct stages defined in the visual system.

### Spontaneous activity reflects functional organization of the developing inferior colliculus

Tonotopy is the characteristic topographic organization of the auditory system. Recent imaging studies directly visualize tonotopic modules in auditory centers (Issa et al., 2014, Wong and Borst, 2019, Romero et al., 2020). In the inferior colliculus, dorsal cortex (DCIC) and lateral cortex (LCIC) display reverse tonotopic gradients at the dorsal surface, with low-frequency zones in the medial and high-frequency zones in the lateral parts (Wong and Borst, 2019, Babola et al., 2018). We resolved this mirror-symmetrical structure from prehearing spontaneous activity with dimensionality reduction techniques (Supplementary Figure 3D), suggesting the existence of dual functional domains before hearing onset. On the other hand, anatomically separate fibro-dendritic laminae (Morest and Oliver, 1984) and a functionally discontinuous tonotopic organization (Malmierca et al., 2008) have been documented in the IC. We observed discrete, but dynamically evolving functional clusters in the IC based on the similarity of activity patterns between individual pixels (Figure 3E). The optimal number of spontaneous activity clusters increased from 2-3 at P0-P1 to 5-6 at ~P12 in each hemisphere. This is consistent with the fact that spontaneous bands became more optically confined at later stages (Figure 1F). Intriguingly, peripheral drive from the cochleae does not become more spatially constrained as animals mature (Tritsch and Bergles, 2010). Therefore, our results suggest a significant refinement of functional circuits in the central auditory system before hearing onset.

---

**Supplementary Figure 1.**
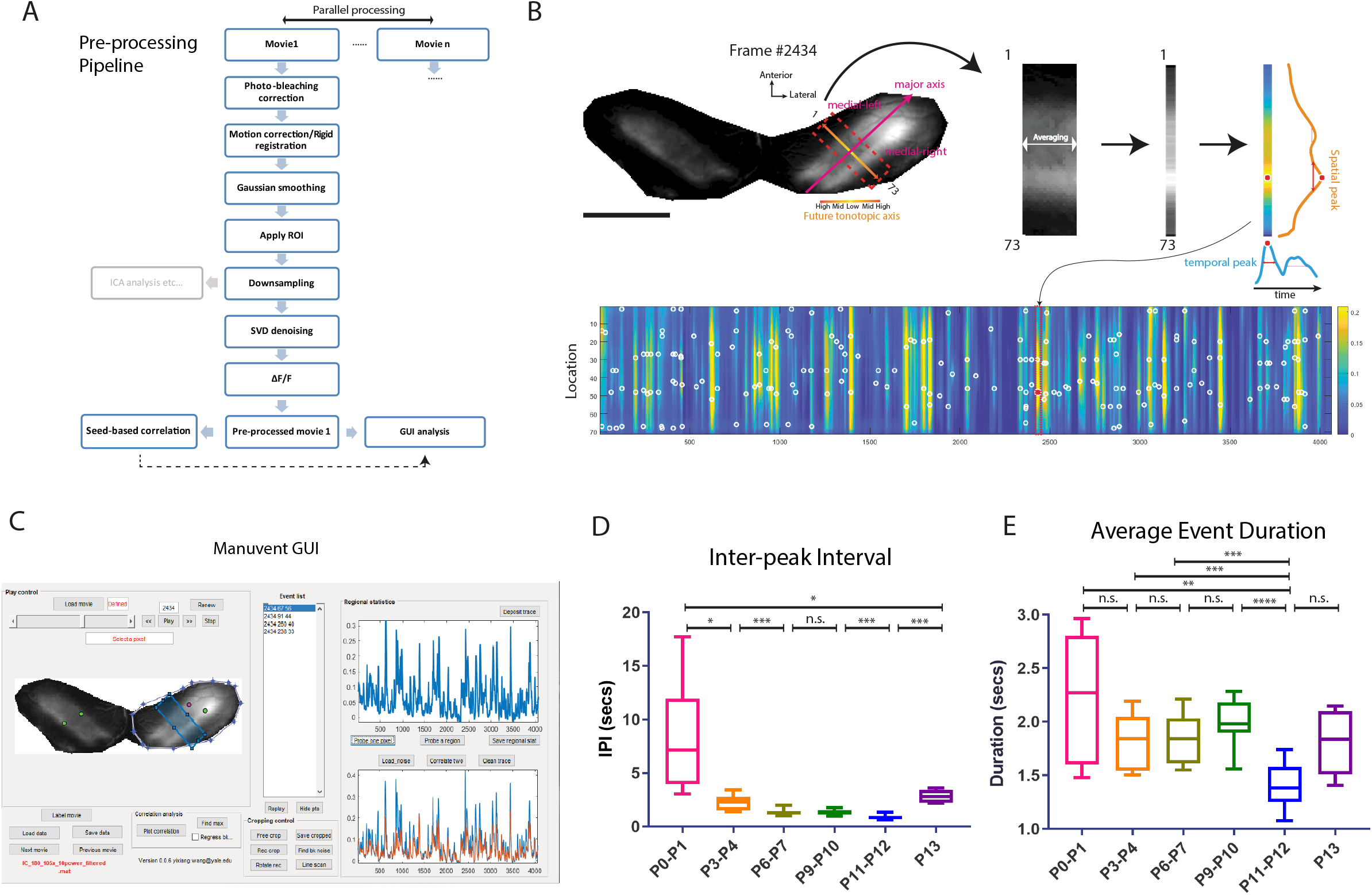
Graphic user interfaces for wide-field imaging data analysis (related to Fig. 1) (A) Object-oriented pre-processing pipeline for high throughput parallel processing (B) Major steps to generate “line-scans” across the future tonotopic axis. Top panels: Left-most image shows an example frame from a continuous movie with an ROI mask over the IC. Magenta arrow represents the major axis of spontaneous bands’ intensity profile. Medial-left and Medial-right regions represent two future tonotopic domains on the opposite sides of the major axis. Yellow to orange gradient denotes the future tonotopic axis/reversal. Dashed red line indicates the rectangular ROI defined for averaging. Averaged one-dimensional line scan is shown in gray and parula color scales. Red dot denotes the intensity peak identified in the line-scan. Orange curve: spatial intensity distribution. Blue curve: temporal intensity trace. Double-headed red arrows on the curves represent half widths of spatial or temporal peaks. Bottom panel: a series of “line-scans” representing a continuous movie. The line scan on the top panel is enclosed in a dashed rectangle. The red dot denotes the same peak as shown above. All other peaks were labeled as empty with circles. Colormap: parula (MATLAB). Scale bar indicates 1 cm. 1 and 73 denotes the first and last pixels. See also STAR Methods. (C) Snapshot of “Manuvent” GUI. Showing the same example movie as in Figure 1. (D) Average inter-peak interval in seconds across age groups. Defined as the mean temporal interval between neighboring peaks in the line-scan analysis. (E) Average event duration across age groups. Defined as the mean temporal half width of all fluorescent peaks in the line-scan analysis. For all Box plots in this study: hinges: 25 percentile (top), 75 percentile (bottom). Box whiskers (bars): Max value (top), Min value (bottom). The line in the middle of the box is plotted at the median. Significance marks: n.s. p>0.05, * p < 0.05, ** p < 0.01, *** p < 0.001, **** p < 0.0001, two-tailed unpaired t test with Welch’s correction. Number of animals: P0-P1 (N = 8); P3-P4 (N = 11); P6-P7 (N = 10); P9-P10 (N = 9); P11-P12 (N = 11); P13 (N =5). Same animals as in Figure 1.

**Supplementary Figure 2.**
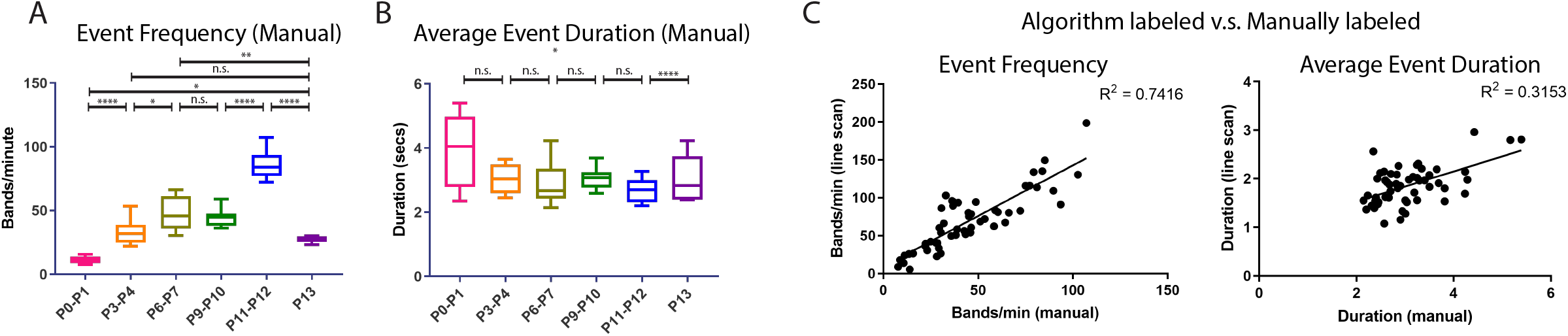
Manually labeled data (related to Fig. 1) (A) Event frequency across age groups (manually labeled). Measure by number of spontaneous bands/events observed per minute in both hemispheres of the IC by a human tester. (B) Average event duration across age groups (manually labeled). Duration is defined as the number of frames in between the first and last frames of events divided by acquisition frequency (10Hz). (C) Linear plots comparing algorithm-generated line-scan results (y axis) and manually labeled results (x axis). Data points include all movies used in Figure 1. Number of animals: P0-P1 (N = 8); P3-P4 (N = 11); P6-P7 (N = 10); P9-P10 (N = 9); P11-P12 (N = 11); P13 (N =5). Same animals as in Figure 1.

**Supplementary Figure 3.**
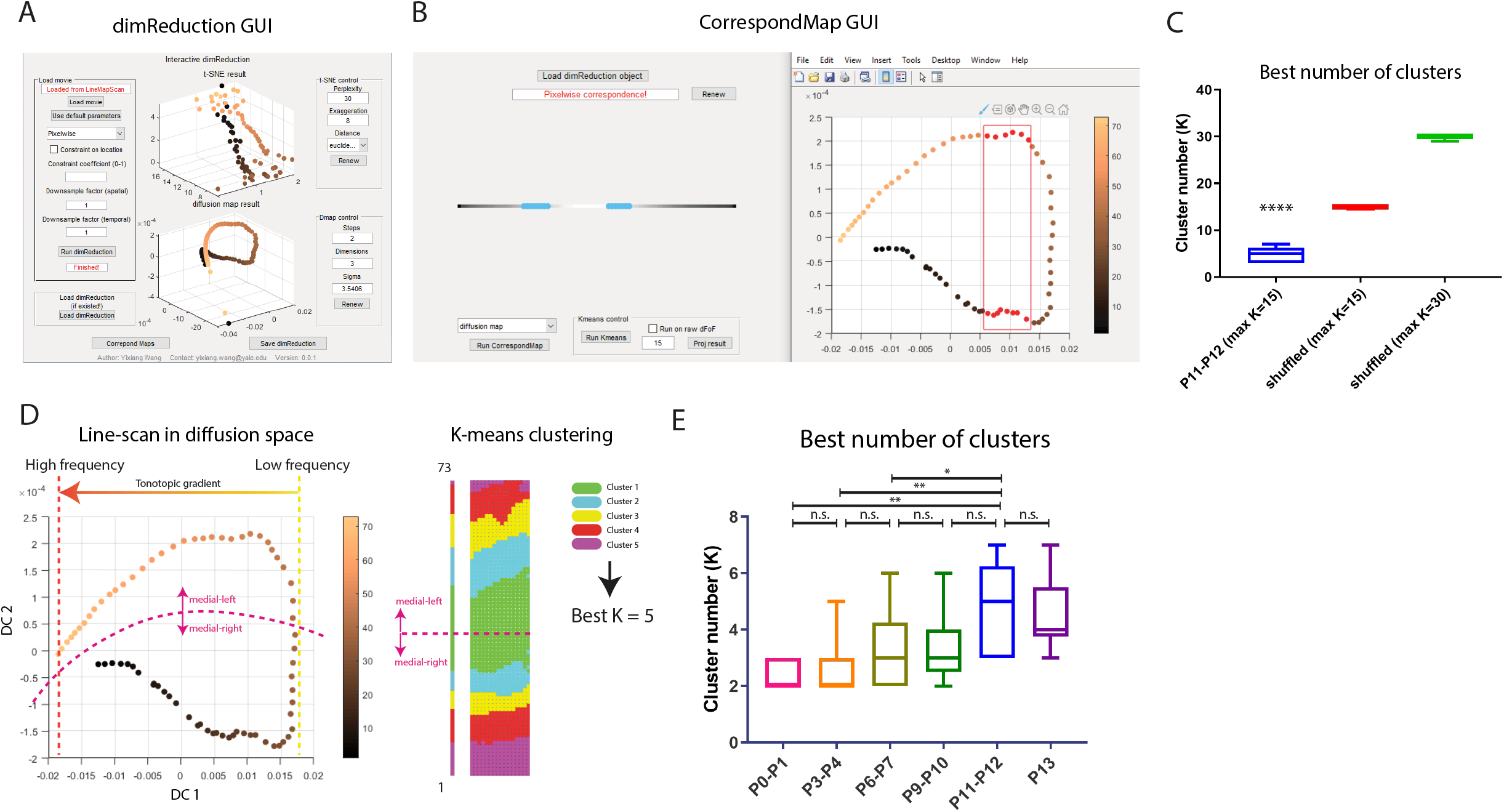
Dimensionality reduction resolves functional organization of the IC (related to Figure 1) (A) Snapshot of “dimReduction” GUI. Showing the same dimensionality reduction result in Figure 1G but with three diffusion components. Also showing dimensionality reduction results based on t-SNE. See also STAR Methods. (B) Snapshot of “CorrespondMap” GUI. Showing the same diffusion map result in Figure 1G. Selected points (red) in the right panel correspond to pixels on the line-scan map in the left panel (light blue). See also STAR Methods. (C) Best number of clusters. Blue: original data at P11-P12. Red: shuffled data with largest possible K equals to 15. Green: shuffled data with largest possible K equals to 30. Number of animals: 11. (D) Left panel: example diffusion map result. Pixels are projected to points in the diffusion space. DC 1/ DC 2: diffusion component 1/2. Single-headed yellow-to-orange arrow denotes the putative future tonotopic gradient. Dashed magenta curve delineates the boundary between the two symmetrical tonotopic domains: medial-left and medial-right. Colorbar denotes pixel number. Colormap: copper (MATLAB). Right panel: example K-means clustering result. Correspond to top panels in Supplementary Figure 1B. (E) Best number of clusters across age groups. Defined as the optimal cluster number evaluated by the Davies-Bouldin criterion. Number of animals: P0-P1 (N = 8); P3-P4 (N = 11); P6-P7 (N = 10); P9-P10 (N = 9); P11-P12 (N = 11); P13 (N =5). Same animals as in Figure 1.

Text for Supplementary Figure 3:

We were also interested in whether the intrinsic dynamics of the spontaneous activity were able to reflect distinct functional organizations of the IC observed in mature animals. To address this question, we took advantage of dimensionality reduction and unsupervised learning techniques. High-dimension calcium imaging data was embedded to low-dimensional manifolds using diffusion map. Pixels on the line-scans differentiated as two mirroring domains along the vertical direction in the diffusion space (divided by a dashed magenta curve, left panel of Supplementary Figure 1D). When slicing the data points horizontally, we were able to extract symmetrically distributed pixels of similar future characteristic frequencies (Supplementary Figure 3B). This is consistent with the reverse organization of dual tonotopic maps in the DCIC and LCIC (Wong and Borst, 2019). We then conducted K-means clustering on pixels given a wide range of possible cluster number/K. We found that pixels indeed formed discrete clusters and the optimal K increased as animals grew older, which peaked at around 5-6 in one hemisphere at P11-P12 (Supplementary Figure 1E). This means that more functional modules emerged as animals matured, consistent with a more refined spatial profile at later stages (Figure 1F). In contrast, shuffled data always achieved optimal conditions close to the maximum possible K, indicating lack of discrete structure (Supplementary Figure 3C). The distribution of functional clusters was mirror-symmetrical (right panel of Supplementary Figure 1D), similar to the diffusion map result. As the IC has anatomically separate fibro-dendritic laminae and functionally discontinuous stepwise response curves (Jeffery A. Winer, 2005, Malmierca et al., 2008), our results indicated that evolving but distinct functional modules existed even before hearing onset.

**Supplementary Figure 4.**
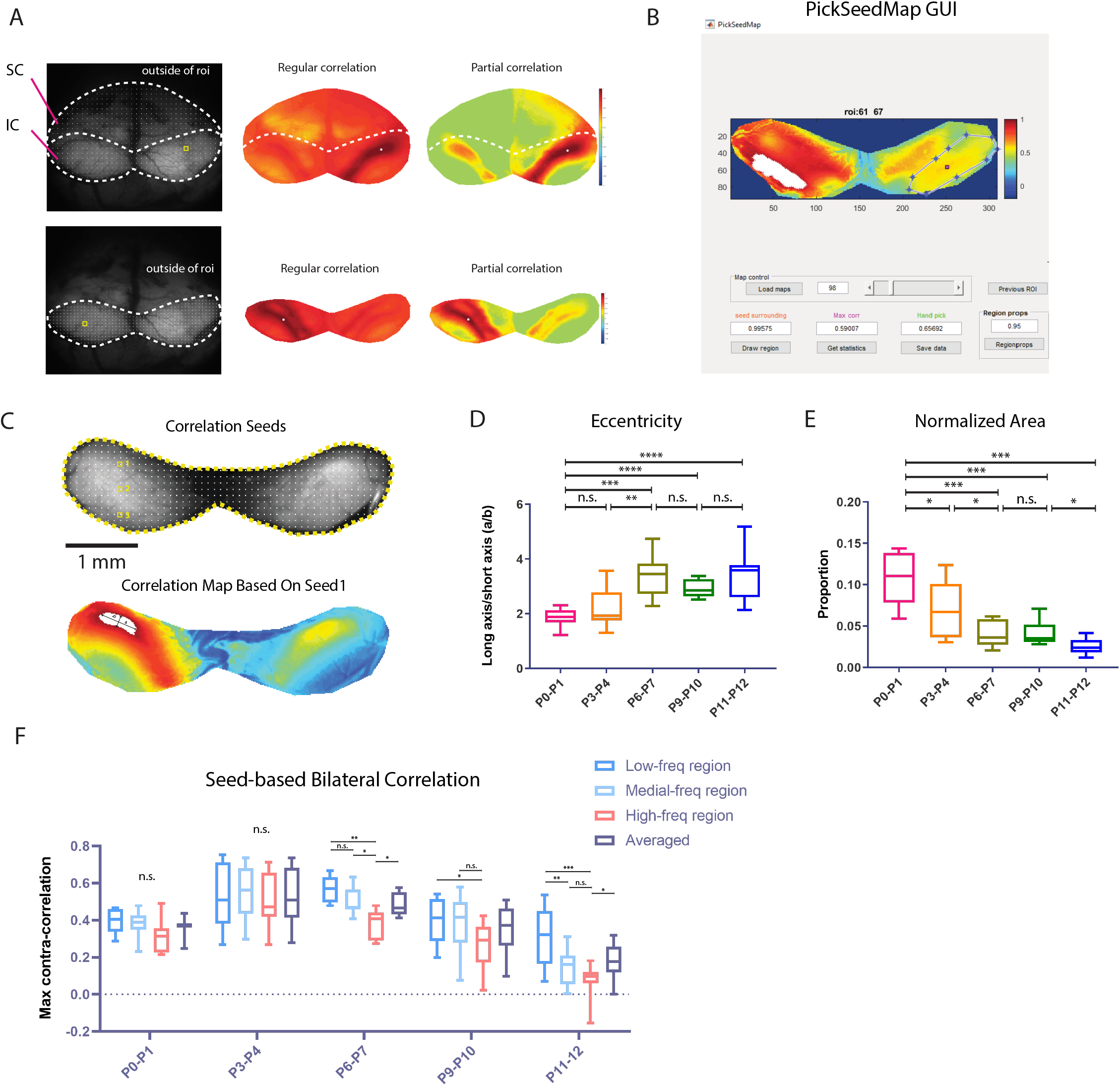
Demonstration of partial correlation (related to Figure 1) (A) Top left panel: Example correlation seeds covering the whole midbrain. Dashed white line delineates the boundary of the ROI mask. Each white dot inside the ROI denotes a reference seed. SC: superior colliculi. IC: inferior colliculi. The seed used for the example correlation maps is labeled with an empty yellow square; Top middle panel: correlation map generated with normal Pearson correlations; Top right panel: correlation map generated with partial correlations controlling on mean activity from pixels outside the ROI. Bottom panels: similar as the top panels but with a ROI mask over the IC only. Note that the color scale here is different from the color scale of all correlation maps in main figures. Colormap: jet (MATLAB). Colorscale: [−1, 1]. See also STAR Methods. (B) Snapshot of “PickSeedMap” GUI. Used for correlation seeds selection and max correlation quantification in the contralateral hemisphere. (C) Top panel: Example of correlation seeds covering the whole IC. Each white dot denotes a reference seed. Three representative seeds used to generate representative correlation maps in Figure 1J are labeled with empty yellow squares and numbers 1,2,3. Bottom panel: a correlation map based on seed number 1. The white region denotes the binary connected component used for ellipse fitting with major (a) and minor (b) axes shown as black double-headed arrows. Colormap: jet (MATLAB). Colorbar: [−0.2, 1]. (D) Eccentricity of fitted ellipses. Defined as the ratio of long axis (a) and short axis (b) (E) Area of fitted ellipses normalized by the size of hemispheres. (F) Figure 1M regrouped by ages to illustrate differences among regions within same age groups. Number of animals: P0-P1 (N = 8); P3-P4 (N = 11); P6-P7 (N = 10); P9-P10 (N = 9); P11-P12 (N = 11). Same animals as in Figure 1.

**Supplementary Figure 5.**
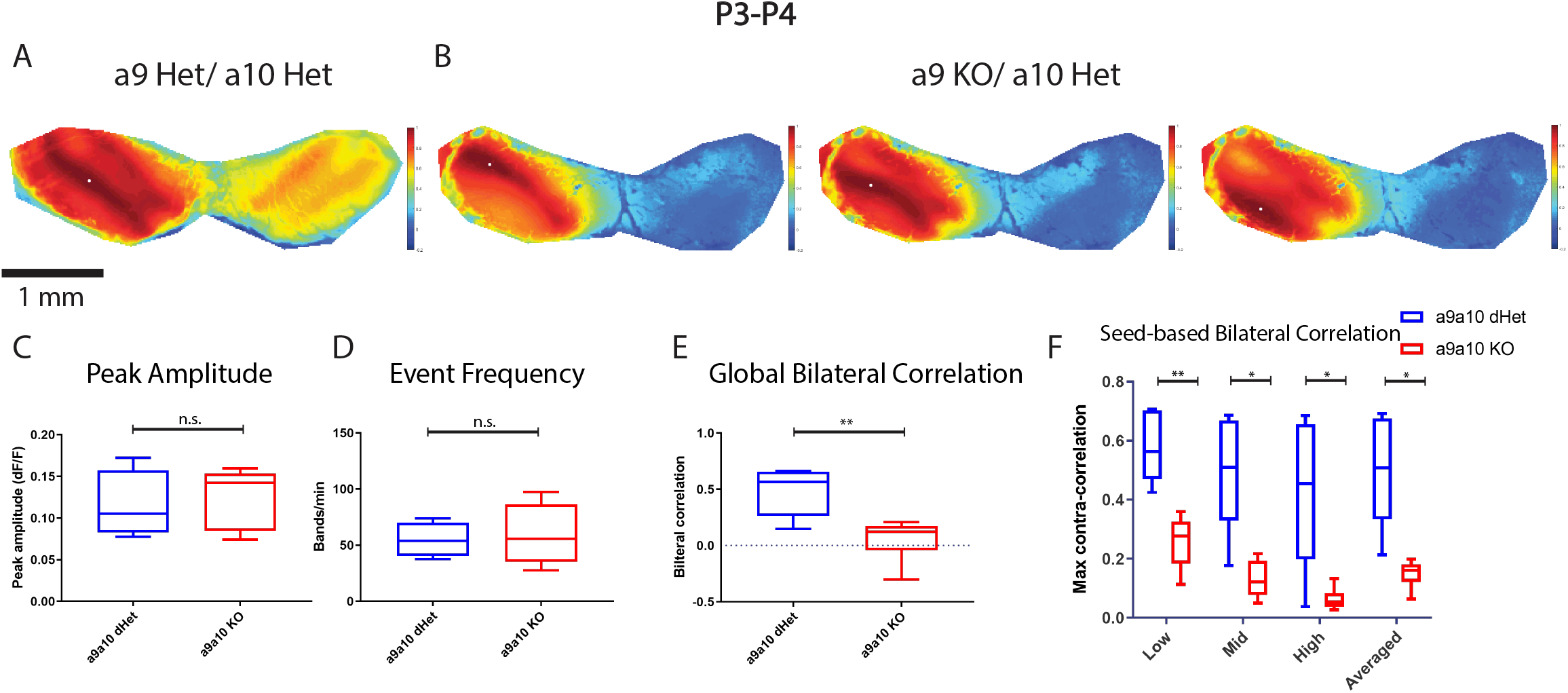
a9a10 nAChR knockouts lack bilateral coupling of spontaneous activity at P3-P4 (Related to Figure 2) (A) Example correlation maps showing correlation patterns in an a9a10 double heterozygous at P3. (B) Example correlation maps showing correlation patterns in a a9 KO a10 Het littermate at P4. (C) Average peak amplitude. Similar to Figure 1D. (D) Event frequency. Similar to Figure 1E. (E) Global bilateral correlation. Similar to Figure 1J. (F) Seed-based bilateral correlation. Similar to Figure 1K. Number of animals: a9a10 dHet (N = 5); a9a10 KO (N = 6).

**Supplementary Figure 6.**
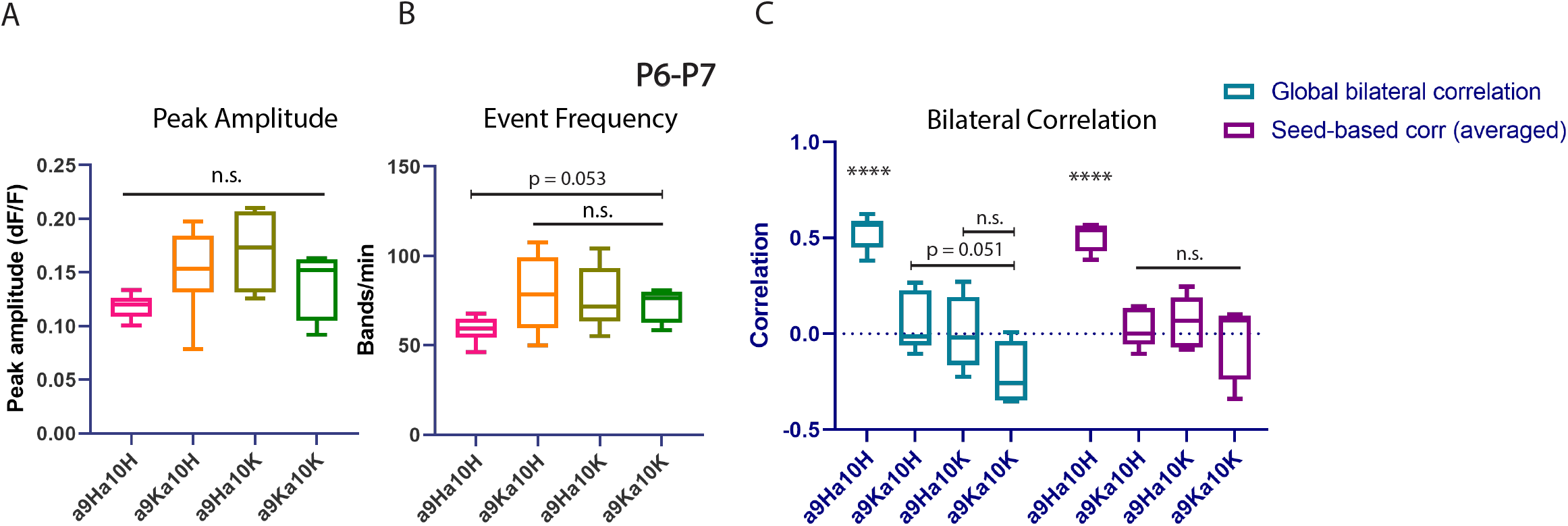
Spatiotemporal and correlation features of a9a10 nAChR single knockouts compared with double knockouts (Related to Figure 2) (A) Average peak amplitude. Similar to Figure 1D. (B) Global and seed-based bilateral correlation of different genotype groups. “Averaged” seed-based correlation is defined as the mean correlation averaged over the three regions. H: heterozygous; K: knockout. For instance, a9Ha10K indicates the a9 heterozygous a10 knockout. Number of animals: a9Ha10H (N = 7); a9Ka10H (N = 5); a9Ha10K (N =5); a9Ka10K (N = 4). All at P6-P7.

**Supplementary Figure 7.**
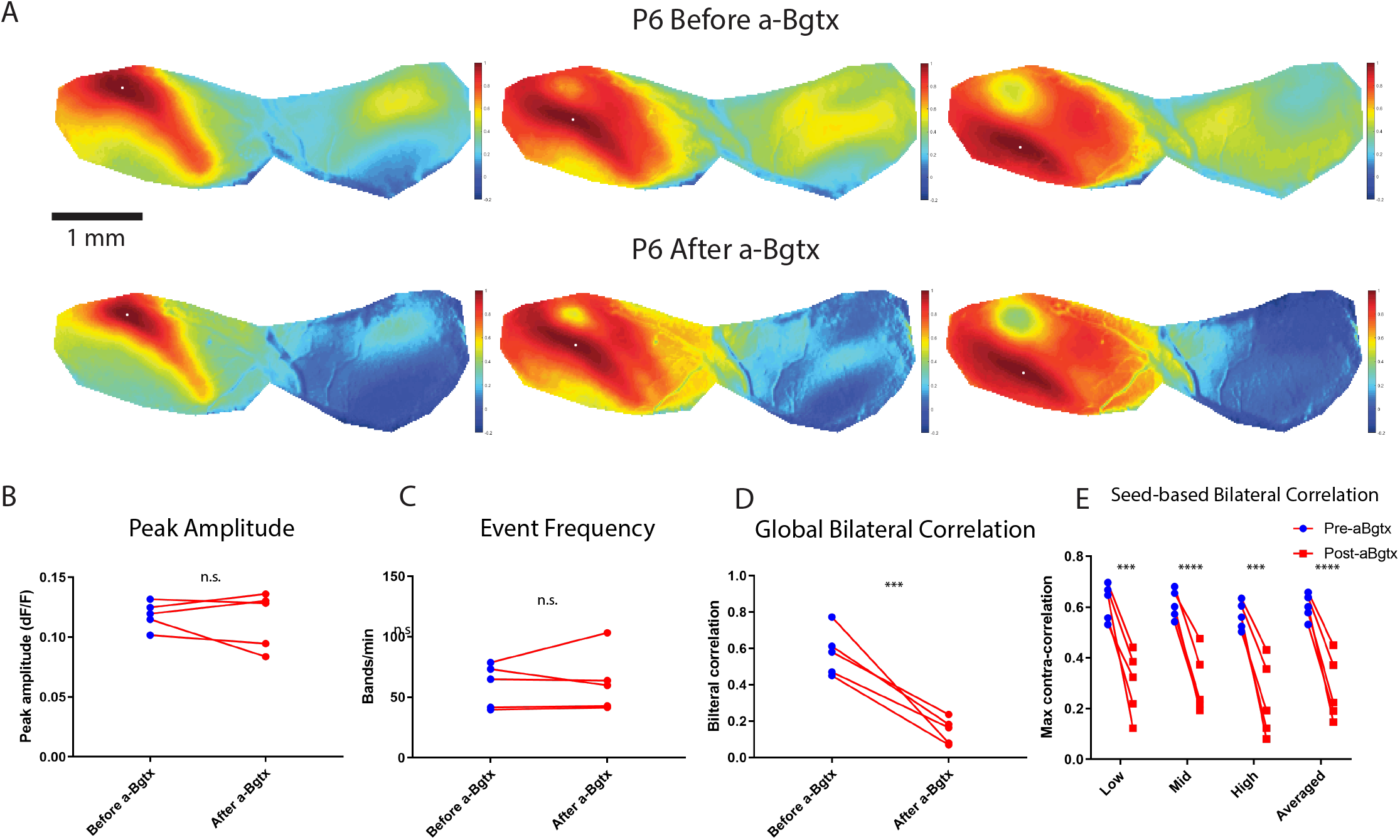
Acute alpha-bungarotoxin application abolishes bilateral coupling in vivo (Related to Figure 3) (A) Example correlation maps showing correlation patterns in the IC before and after alpha-bungarotoxin from the same SNAP25-G6s animal at P6. Top panels: before apamin. Bottom panels: after apamin. Three typical representative seeds (white dots) located in the future low-, mid-, and high-frequency regions. Similar to Figure 3E. (B) Average peak amplitude. Similar to Figure 1D. (C) Event frequency. Similar to Figure 1E. (D) Global bilateral correlation. Similar to Figure 1J. (E) Seed-based bilateral correlation. Similar to Figure 1K. Number of animals (SNAP25-G6s) = 5. Scale bar denotes 1 mm.

**Supplementary Figure 8.**
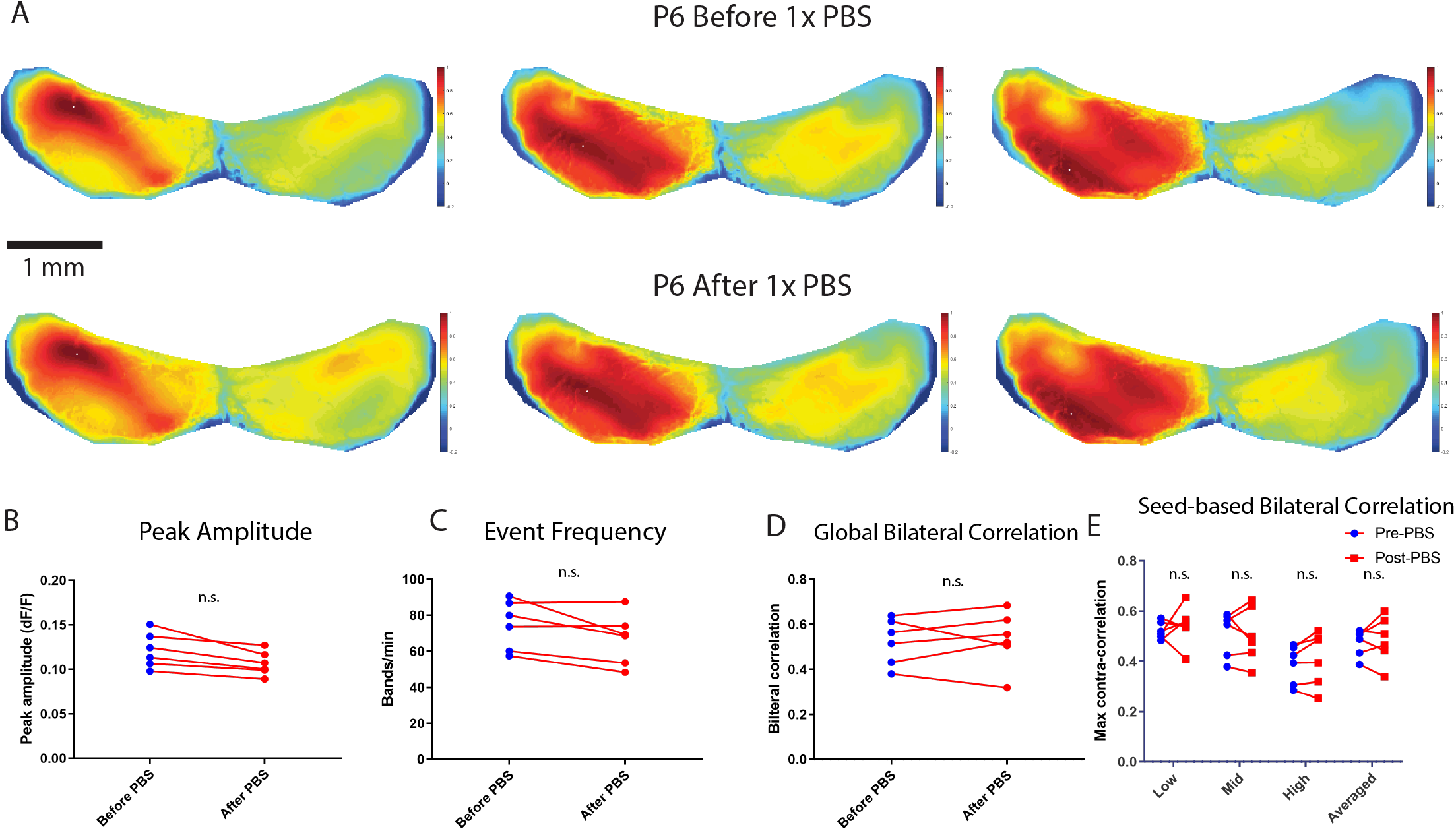
Acute saline application does not affect bilateral coupling in vivo (Related to Figure 3) (A) Example correlation maps showing correlation patterns in the IC before and after 1XPBS from the same SNAP25-G6s animal at P6. Top panels: before saline. Bottom panels: after saline. Three typical representative seeds (white dots) located in the future low-, mid-, and high-frequency regions. Similar to Figure 3E. (B) Average peak amplitude. Similar to Figure 1D. (C) Event frequency. Similar to Figure 1E. (D) Global bilateral correlation. Similar to Figure 1J. (E) Seed-based bilateral correlation. Similar to Figure 1K. Number of animals (SNAP25-G6s) = 5. Scale bar denotes 1 mm.

**Supplementary Figure 9.**
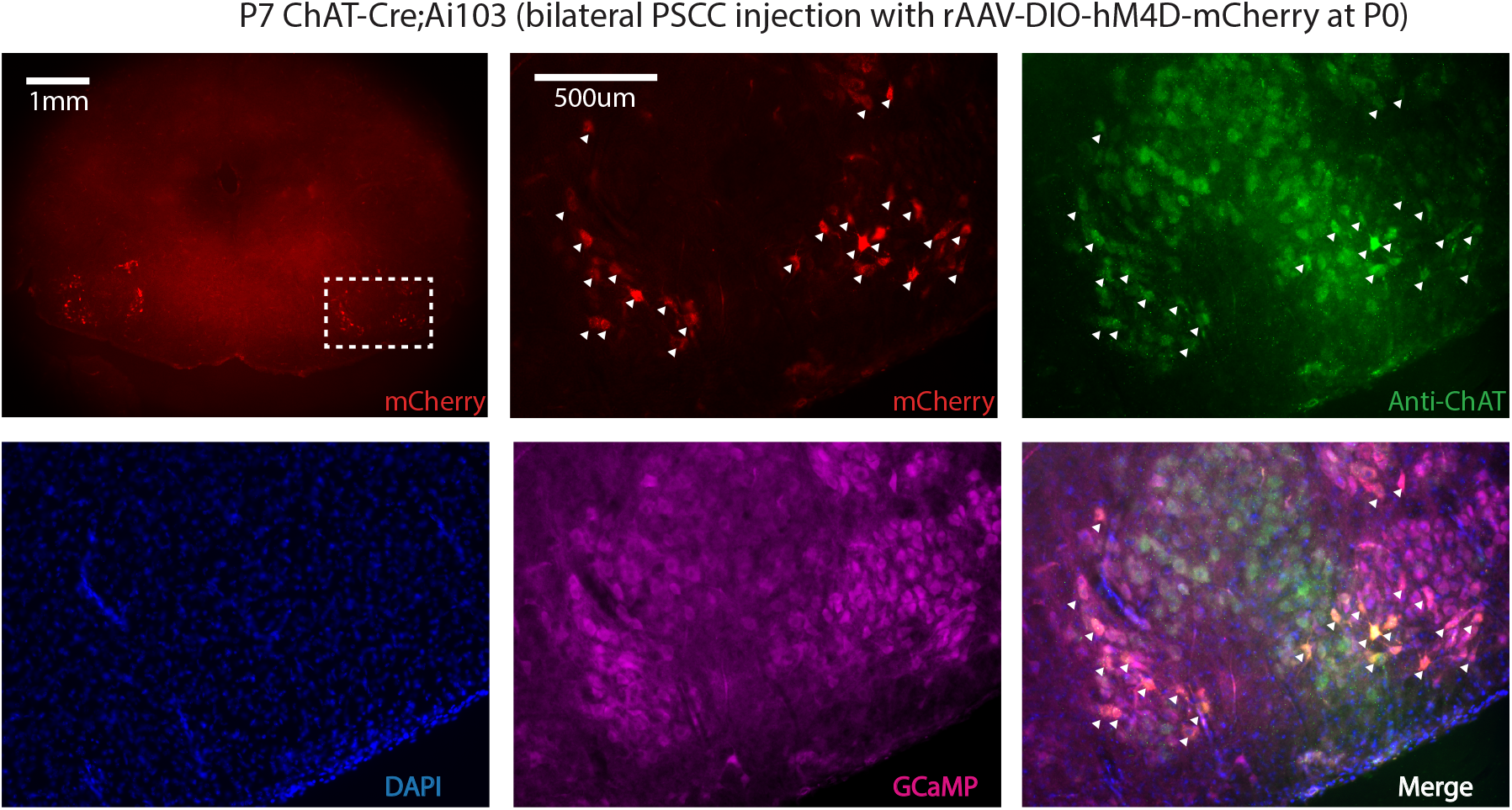
Cholinergic olivocochlear neurons are targeted specifically (related to Fig. 4) First panel: A brainstem section that contains olivocochlear neurons expressing mCherry. Magnification: 2.5X. Scale bar: 1mm. Following panels: high magnification images of the white dashed rectangle in the first panel: showing mCherry, Alexa 647, DAPI, EGFP channels, and the merged image. Magnification: 10x. Scale bar: 500um. Experimental information: ChAT-Cre;Ai103 animal was perfused at P7 (bilateral PSCC injection with rAAV-DIO-hM4D-mCherry at P0). White arrowheads denote cells that express mCherry/infected by the retrograde virus. See also STAR Methods.

**Supplementary Figure 10.**
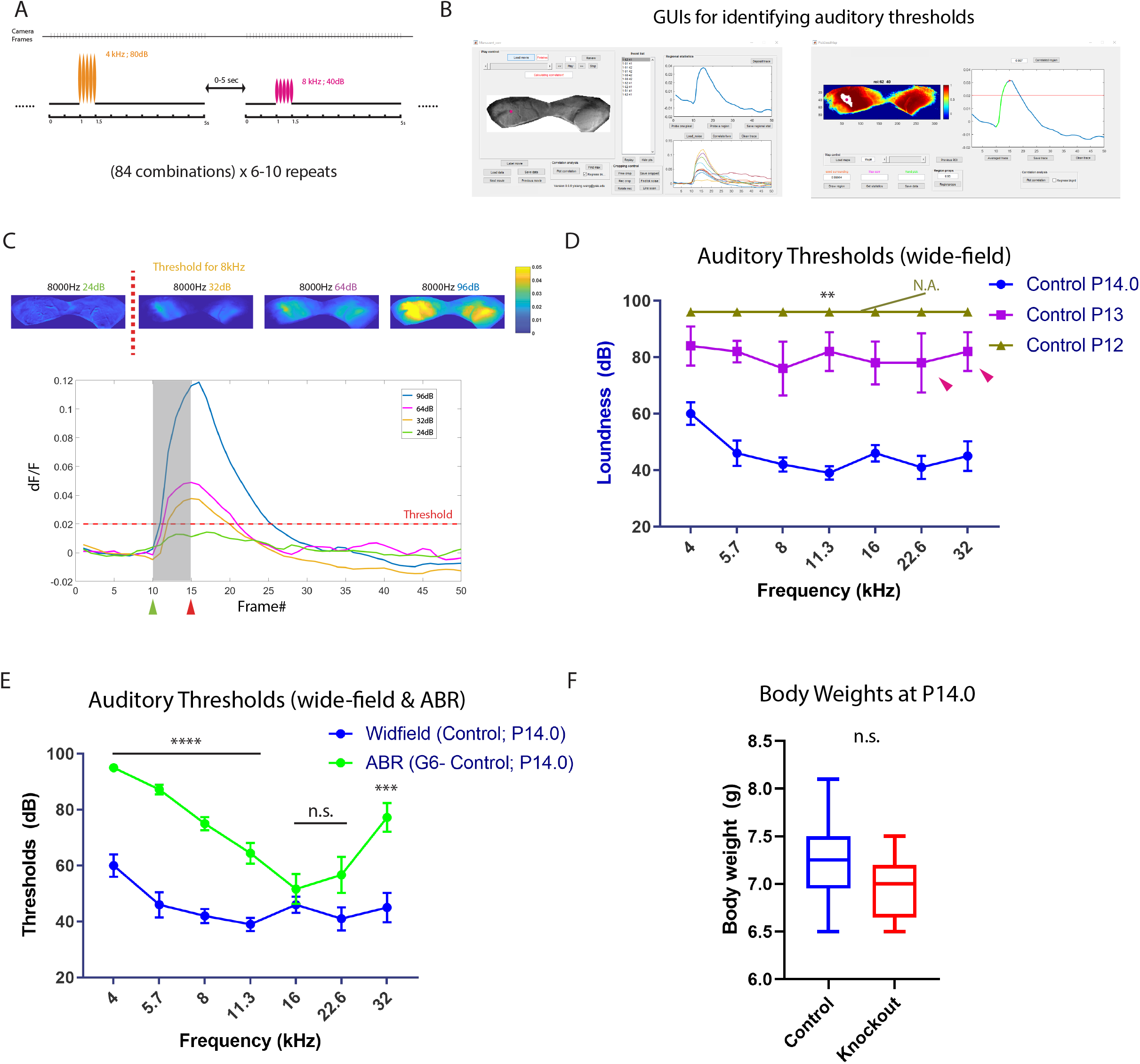
Wide-field calcium imaging resolves auditory thresholds at lower levels (related to Figure 5) (A) Diagram illustrating scheme of acoustic stimuli. Ticks on the “Camera Frames” axis represent endpoints of every frame. Each trial contains 5 seconds of activity/50 frames (acquired at 10 Hz). SAM tones illustrated as wave packets. Note that heights of wave packets are disproportionate to actual voltage amplitudes. See also STAR Methods. Sec/s: second. See also STAR Methods. (B) GUIs “Manuvent” and “PickSeedMap” are compatible with functions to identify auditory thresholds. (C) Top panels: example ΔF/F_0_ responses to 8kHz tones at four different decibel levels. Colormap: parula (MATLAB). Bottom panel: corresponding ΔF/F_0_ curves. Dashed red line indicates threshold. Shaded area denotes the period during which acoustic stimuli are presented. (D) Auditory thresholds measured by wide-field imaging in control animals at postnatal day 12,13, and 14.0. Number of animals: Control P14.0 group = 8 (same animals as in Figure 5E, wide type SNAP25-G6s = 6; a9a10 double heterozygous SNAP25-G6s = 2). Control P13 group = 5 (wide type SNAP25-G6s). Control P12 group = 4 (wide type SNAP25-G6s). N.A. indicates “not available” in the P12 group (no response at the maximum decibel level across the spectrum). Magenta arrowheads indicate that part of the data is not available in the P13 group (1, and 2 animals did not respond at 22.6, and 32 kHz respectively). ** p < 0.01 (note that some p values are smaller than 0.001) two-tailed unpaired t test with Welch’s correction conducted between P13 group and P14.0 group at each frequency. Error bars indicate SEM. (E) Compare auditory thresholds measured by wide-field imaging and auditory brainstem response recording in control animals at P14.0. Number of animals: Widefield P14.0 group = 8 (same animals as in Figure 5E, wide type SNAP25-G6s = 6; a9a10 double heterozygous SNAP25-G6s = 2). ABR P14.0 group = 9 (GCaMP6s negative littermates, same animals as in Figure 5B). Magenta arrowheads indicate that part of the data is not available in the ABR recording. n.s. p>0.05, ***p<0.001, **** p < 0.0001, two-tailed unpaired t test with Welch’s correction conducted between two conditions at each frequency using available data. Error bars indicate SEM. (F) Body weights in grams. Control P14.0 group = 8 (same animals as in Figure 5E, wide type SNAP25-G6s = 6; a9a10 double heterozygous SNAP25-G6s = 2). Knockout = 8 (mix of single knockout of either a9 or a10 subunit and double knockout). Box plot hinges: 25 percentile (top), 75 percentile (bottom). Box whiskers (bars): Max value (top), Min value (bottom). The line in the middle of the box is plotted at the median. Significance marks: n.s. p>0.05, two-tailed unpaired t test with Welch’s correction.

**Supplementary Figure 11.**
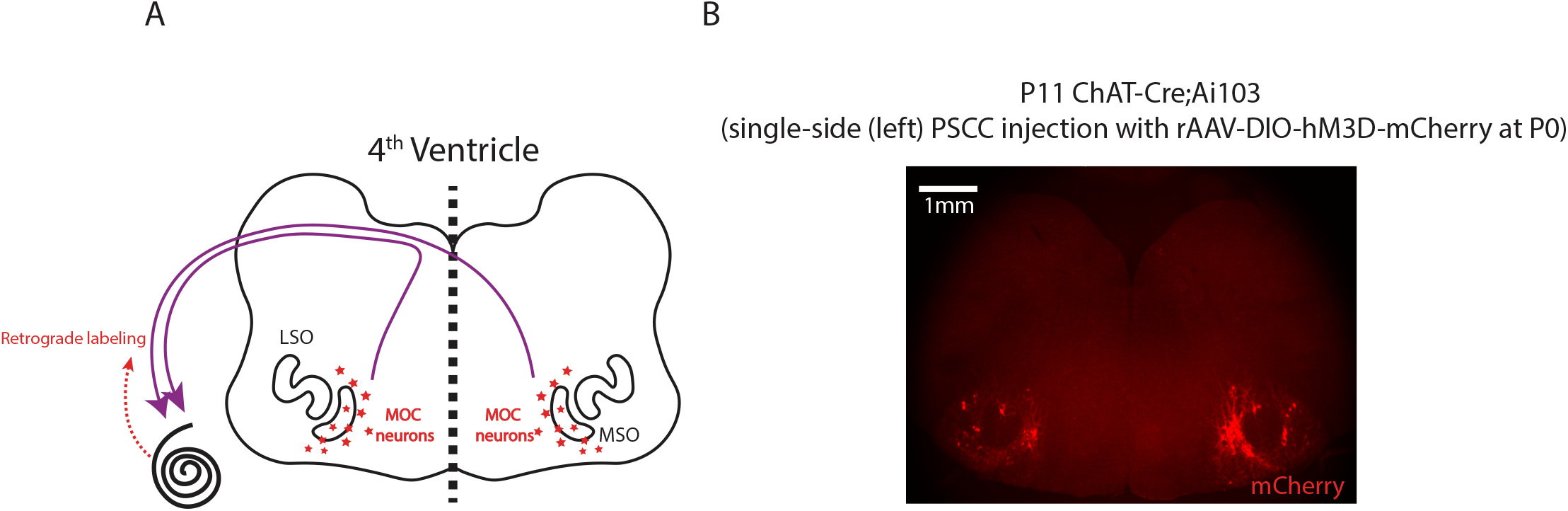
Cochleae received bilateral MOC efferent feedback (related to Figure 6) (A) Schematics of cochlea receiving bilateral efferent feedback from the medial olivocochlear neurons (MOC). LSO: lateral superior olive; MSO: medial superior olive. (B) A brainstem section that contains olivocochlear neurons expressing mCherry. Magnification: 2.5X. Scale bar: 1mm. Experimental information: ChAT-Cre;Ai103 animal was perfused at P11 (single-side PSCC injection to the left cochlea with rAAV-DIO-hM3D-mCherry at P0).

## Acknowledgments

We would like to thank all members of the Crair lab, the Santos-Sacchi lab, and the Navaratnam lab for their helpful comments on this project. We would like to thank Dr. Leonard Kaczmarek and Dr. Yalan Zhang for providing apamin for pilot pharmacological experiments. We would like to thank Dr. Winston Tan and Dr. Jun-ping Bai for providing guidance on conducting ABR experiments. We would like to thank Dr. Steven W. Zucker and for suggestions on applying diffusion map analysis. We would like to thank Dr. Rui Chang and Chuyue Yu for providing Alexa Fluor® 647 AffiniPure Donkey Anti-Goat IgG (H+L) anitbody. We would like to thank Dr. Jessica Cardin on discussions regarding this study. Also thanks the family of William Ziegler III for their support.

## Author Contributions

Conceptualization, Y.W., A.G. and M.C.C.; Methodology, Y.W., A.G., L.S., and M.C.C.; Software, Y.W.; Formal Analysis, Y.W., and M.S.; Investigation, Y.W.; Resources, B.M., J.S., D.N., and M.C.C.; Writing– Original Draft, Y.W.; Writing–Review & Editing, Y.W., M.S., L.S., A.G., and M.C.C.; Visualization, Y.W.; Supervision, M.C.C; Project Administration, Y.W., Y.Z., M.C.C.

## Declaration of Interests

The authors declare no competing interests.

## Star Methods

### Lead Contact and Materials Availability

Further information and requests for resources and reagents should be directed to and will be fulfilled by the Lead Contact, Michael C. Crair. This study did not generate new unique reagents.

### Experimental Model and Subject Details

#### Transgenic Models

##### SNAP25-G6s animals

B6.Cg-Snap25tm3.1Hze/J (JAX#025111) animals that have pan-neuronal GCaMP6s expression were used widely in this study.

##### a9/a10;SNAP25-G6s animals

a9/a10 nAChR double-knockout (a9^−/−^a10^−/−^) animals on C57BL/6 background were obtained from Dr. Barbara Morley (Morley et al., 2017). a9^−/−^a10^−/−^ line was crossed to SNAP25-G6s line (JAX #025111) to generate double-heterozygous animals that express GCaMP6s (a9^+/−^ a10^+/−^; SNAP25-G6s^+/null^). These animals were backcrossed to a a9^−/−^a10^−/−^ line to generate four different genotypes of offspring (double-heterozygous: a9^+/−^a10^+/−^, single-knockout: a9^+/−^a10^−/−^, a9^−/−^a10^+/−^, double-knockout: a9^−/−^a10^−/−^) with or without GCaMP6s expression (SNAP25-G6s^+/null^ or SNAP25-G6s^null/null^).

##### ChAT-Cre;SNAP25-G6s animals

Homozygous ChAT-Cre^+/+^ animals (JAX #018957) were crossed to SNAP25-G6s animal (JAX #025111) to generate ChAT-Cre heterozygous offspring that express GCaMP6s (ChAT-Cre^+/−^; SNAP25-G6s^+/null^).

#### Animals Usage

Animals of both sexes were used in this study. Animal care and use followed the Yale Institutional Animal Care and Use Committee (IACUC), the US Department of Health and the Human Services, and institution guidelines. In Figures 1, SNAP25-G6s animals between P0-P13 were used for spatiotemporal and correlation analysis at different ages. In Figure 2 and related Supplementary Figures 5-6, GCaMP6s-positive a9/a10;SNAP25-G6s animals (see Transgenic Models) were used for spatiotemporal and correlation analysis at P6-P7 (Figure 2 and Supplementary Figure 6) or at P3-4 (Supplementary Figure 5). In Figures 3 and related Supplementary Figures 7-8, SNAP25-G6s animals between P5-7 were used for in vivo pharmacological experiments. In Figure 4, GCaMP6s-positive ChAT-Cre;SNAP25-G6s animals (see Transgenic Models) were used for chemogenetic experiments. In Figure 5 and the related Supplementary Figure 10, the control group consists of wide-type SNAP25-G6s animals and a9/a10 double heterozygous animals. The a9/a10 knockouts consists of single and double knockout of a9 and/or a10 subunits. GCaMP6s positive animals were used for auditory thresholds measurement with wide-field imaging. Their GCaMP6s negative littermates were used for auditory brainstem response.

#### Surgery For In Vivo Imaging

##### Installation of cranial windows

Mice were installed with cranial windows using procedures similar to (Ackman et al., 2012) with modifications: 1. All animals were head-fixed to an articulating base stage (SL20, Thorlabs) with optic posts and angle post clamps that allowed rapid angular positioning 2. To expose the entire dorsal surface of inferior colliculi (IC), the skull over IC and anterior-dorsal part of cerebellum was removed. 3. For animals younger than P4, initial isoflurane concentration for anesthesia was adjusted to 2% instead of 2.5%. 4. As spontaneous activity started to recover ~30 mins after anesthesia (Ackman et al., 2012), all animals were allowed at least one hour to recover (with oxygen delivered) before imaging sessions. 5. Animals were nested with cotton gauze as previously described (Gribizis et al., 2019). 6. For experiments involving bilateral A1 and IC (Figures 2-3), the skull over A1 was rinsed with 1x PBS and cleaned with PVA eye spears. A minimal amount of cyanoacrylate glue was carefully applied to the cleaned skull to create a smooth surface for direct imaging through the skull.

##### In vivo pharmacology via round window

After acquisition of non-manipulated data, animals were anesthetized with 2.5% isoflurane again. Articulating stage was tilted ~45° and angle post clamps were adjusted to allow better angular positions to perform postauricular incisions. Procedures to expose round window niches are similar to (Babola et al., 2018). To facilitate robust pharmacological delivery, round window membrane was punctured and removed using fine forceps. Fluid efflux was drained with sterile filter paper. Gelform soaked with 1uL of different pharmacological compounds (apamin: 200uM; alpha-bungarotoxin: 3mM; or 1x PBS) was shallowly inserted into the cochlea through the round window. The opening was quickly sealed with cyanoacrylate glue using a plastic glue stick. The surgery was performed on both sides of cochleae in ~20 minutes (isoflurane was changed to 1.5% during the surgery and stayed off during recovery). Animals were allowed at least one hour to recover (with oxygen delivered) before imaging sessions.

#### In Vivo Calcium Imaging

Schematics of wide-field imaging apparatus is shown in Figure 1A. The microscope was placed in a soundproof booth as described in Acoustic Stimulation For In vivo Calcium Imaging, this paper.

Procedures for imaging were almost identical to (Babola et al., 2018) with three exceptions: 1. Pups were wrapped with cotton gauze during data acquisition as described in (Gribizis et al., 2019) instead of being placed in a swaddling 15 mL conical centrifuge tube 2. The size of field of view varied across conditions to maximize visibility of regions of interest (IC only or simultaneous imaging over auditory cortex and IC). 3. Each recording session contained continuously acquired movies for more than 30 mins.

#### Chemogenetic Experiments

##### Posterior semicircular canal (PSCC) injection in neonates

Neonatal ChAT-Cre;SNAP25-G6s animals (see Transgenic Models) were genotyped and GCaMP6s-positive pups were selected for PSCC injection at P0-P1. Procedures were similar to (Isgrig and Chien, 2018) with modifications: 1. Pups were placed in crushed ice for ~90 seconds for initial anesthesia 2. Athetized pups were placed on a reusable gel ice pack with crushed ice around the body. 3. 2uL (1 uL each side) of floxed retrograde AAV was injected (rAAV-hSyn-DIO-hM4D-mCherry: titer ≥ 8×10^12^ vg/mL; or rAAV-hSyn-DIO-hM3D-mCherry: titer ≥ 7×10^12^ vg/mL, see Key Resource Table) via micro-injector. All procedures were finished in 10 minutes.

##### Clozapine N-oxide (CNO) injection

CNO dose was 5mg/kg for inhibitory-DREADD (hM4Di) experiments (Fig. 4A-E) and 1mg/kg for excitatory-DREADD (hM3Dq) experiments (Fig. 4F-J). After acquisition of non-manipulated data, a fixed volume of CNO (various concentrations to match body weights with required doses) was injected intraperitoneally (IP injection) to animals. As reported previously (Jendryka et al., 2019), an adequate amount of CNO can be detected in brain tissues 15 mins after injection. Imaging sessions started ~15 mins after IP injection and lasted for 30 mins.

##### Immunohistochemistry

Mice were anesthetized via IP injection with a ‘rodent combination cocktail’ (Ketamine 37.5 mg/ml, Xylazine 1.9 mg/ml and Acepromazine 0.37 mg/ml) at 1ml/kg and perfused transcardially with 1x PBS followed with 4% PFA. Brains were removed and fixed overnight in 4% PFA. After embedding brains in 2.5% agarose (made in 1x PBS), coronal sections (50um) were collected using vibratome (Leica VT1000 S). Brain slices were permeabilized in 0.7% Triton X-100 in 1x PBS for 30 mins before blocking. Permeabilized slices were rinsed in PBST (1x PBS with 0.01% Triton X-100) three times (10 mins each time) and incubated with blocking solution (10% donkey serum, 1% bovine serum albumin, 0.5% Triton X-100 in 1x PBS) for 24 hours at 4 C°. Brain slices were rinsed 3 times in PBST and then incubated with primary antibody (1:400; goat anti-choline acetyltransferase (ChAT), catalog# AB144P, Millipore) for ~48 hours at 4 C°. After primary incubation, the slices were rinsed 3 times in PBST and then incubated with secondary antibody (1:500 Alexa 647-conjugated Donkey Anti-Goat IgG, Cat# 705-605-147, Jackson ImmunoResearch) at room temperature for ~3 hours. After secondary incubation, the slices were rinsed 3 times in PBST and mounted on glass slides with antifade mounting medium with DAPI (VECTASHIELD). Images were acquired using a Zeiss Axio Imager Z2 equipped with a CCD camera (AxioCam HRC, Carl Zeiss). Merged images were constructed using Zeiss ZEN Blue software from four different channels with pseudocolors (Supplementary Figure 9: Blue: DAPI; Magenta: GCaMP; Red: mCherry; Green: Alexa 647).

#### Acoustic Stimulation For In Vivo Calcium Imaging

Wide-field microscope was enclosed in a double-pane booth made of soundproof plywood and acoustic foam (See Figure 5D). Background noise was <30 dB SPL inside the booth (the major source was the CMOS camera fan at the top of the wide-field microscope, measured with a BK Precision 735 sound level meter). An electrostatic speaker (ES1, Tucker-Davis Technologies) was placed 5cm from lambda and parallel to the sagittal suture. Acoustic stimuli were generated with customized open-source software in MATLAB (“Baphy”, Neural Systems Laboratory, University of Maryland College Park). A digital-to-analog converter (National Instruments BNC-2110) transformed MATLAB outputs to voltage signals to the speaker driver (ED1, Tucker-Davis Technologies). To optimize calcium responses in wide-field imaging setting (Issa et al., 2014), sinusoidal amplitude modulated tones (SAM, modulation depth = 1; modulation frequency = 10Hz) were presented free field at frequencies between 4 kHz to 32 kHz (at half-octave step) and sound intensities from 96 dB SPL to 8 dB SPL (in 8 dB decrements). This resulted in 84 different frequency-SPL combinations. Each stimulation trial was 5-seconds long consisting of: 1-second pre-stimulation idle time, 0.5-second SAM acoustic stimulation, and 3.5-seconds post-stimulation idle time. Trials were spaced by random intervals (between 0-5 seconds). Camera frames were triggered by and time-locked with acoustic stimuli via a neurophysiological stimulator (Master 8, A.M.P.I. Israel) so that each stimulation trial rendered exactly 50 camera frames (movies were acquired at 10 Hz). One stimulation session contained all 80 frequency-SPL combinations in random order and produced a 4000-frame (400-second) movie. Digital signals from MATLAB, analog signals (voltage inputs) to the speaker driver, and camera feedback were recorded in a multi-channel data acquisition platform (SPIKE2, CED) at 125kHz sampling rate for quality control and data analysis. An ultrasound microphone was placed 5cm away from the ES1 speaker to measure SPL for response curve calibration (Knowles FG, Avisoft Bioacoustics). Sound levels (pure tones) produced by the ES1 speaker were calibrated to be consistent with the outputs of the FF1 speaker described in the ABR section (±3 dB error). Each animal was presented with 6-10 stimulation sessions in total of 1-2 hours.

To pinpoint birth time accurately, cages were checked twice a day (~12 hours apart). Acoustic stimulation experiments were conducted at P14.0 (within 12 hours interval from 13*24hours to 13*24+12hours after birth), P13 (within 24 hours interval from 12*24 to 13*24 hours after birth) or P12 (within 24 hours interval from 11*24 to 12*24 hours after birth).

#### Auditory Brainstem Response (ABR)

ABR experiments in this study were conducted using the same apparatus (TDT3 system with a FF1 speaker, Tucker-Davis Technologies) and procedures described in (Tan et al., 2017, El-Hassar et al., 2019). Auditory thresholds were determined based on waves I and II (see also Figure 5A). Genotype information was blinded to the tester. In brief, differentially recorded scalp potentials were bandpass filtered between 0.05 and 3 kHz over a 15-ms epoch. A total of 400 trials were averaged for each waveform for each stimulus condition. Symmetrically shaped tone bursts were 3ms long (1 ms raised cosine on/off ramps and 1ms plateau) and were delivered at a rate of approximately 20 per second. Stimuli were presented at frequencies between 2 and 32 kHz and in 5 dB decrements of sound intensity from 105 dB SPL. The ABR threshold was defined as the lowest intensity (to the nearest 5 dB) capable of evoking a reproducible, visually detectable response.

### Data Analysis

#### Image processing

##### Pre-processing

Raw TIFF movies were pre-processed with an object-oriented pipeline. Parallel computing was realized in MATLAB on Yale’s high-performance clusters (Yale Center for Research Computing). The pipeline included following steps (see Supplementary Figure 1A): 1. Photobleaching correction using a single-term exponential fit on averaged fluorescent intensity (frame-wise). 2. Motion correction/subpixel rigid registration (Guizar-Sicairos et al., 2008). 3. Gaussian smoothing (sigma/filter size = 1). 4. ROI mask application (manually defined in ImageJ). 5. Downsampling (average-pooling with a 2×2 filter). 6. Denoising based on singular-vector decomposition 7. Intensity normalization (ΔF/F0, where F0 was defined as the 5^th^ percentile value for each pixel). Key parameters can be customized and were kept consistent across conditions.

##### Seed-based correlation analysis

Number of reference seeds was pre-defined before parallel processing. Movies with a down-sampled field of view were densely covered by 1000 evenly spaced seeds. Regular or partial Pearson-correlation matrices were generated with seeds inside ROI masks using built-in MATLAB functions “corr” and “partialcorr” respectively (see Supplementary Figure 4). For partial-correlation matrices, correlations between reference seeds and other pixels in the ROI were controlled for the averaged fluorescence trace over all pixels outside the ROI (to regress out the influence of non-specific whole-brain fluctuations). All seed-based correlation analysis in this study is based on partial Pearson-correlation except for the demonstration in Supplementary Figure 4A. All seed-based correlation maps were manifested with jet (256) colormap and [-0.2, 1] color scale except for the demonstration in Supplementary Figure 4A (with [-0.1, 1] color scale to better illustration near-zero correlations). Frames identified as containing motions (when amplitudes of subpixel rigid registration were larger than 0.5 pixel) were excluded using motion correction information generated in pre-processing procedures.

#### Graphic-user-interface For Wide-field Auditory Data Analysis

A novel Graphic-user-interface (“Manuvent” GUI, see Supplementary Figure 1C) was designed for interactive data analysis based on pre-processed movies. This multi-purpose platform allowed following analysis:

##### Line scan and peak detection

Rectangular ROIs were interactively drawn in the GUI. Cropped movies were averaged over the direction parallel to the major axis of spontaneous bands to reduce two-dimensional frames to one-dimensional linear representations across the tonotopic axis (see Supplementary Figure 1B, similar to Babola et al., 2018). This process projected a three-dimensional movie (two-dimensional frames × one-dimensional time) to a two-dimensional line-scan map (one-dimensional line representations × one-dimensional time). Spontaneous events were represented as two-dimensional ΔF/F_0_ bumps in the line-scan maps (Supplementary Figure 1B). Peak detection of spontaneous events (finding local maximum of ΔF/F_0_ bumps) was accomplished in an auxiliary GUI (“LineMapScan”, see Supplementary Figure 1B). Line-scan maps were smoothed with a 5×5 spatiotemporal filter (much smaller than minimum spatial and temporal half-width of spontaneous events across ages) before peak detection. 5% ΔF/F_0_ threshold was consistently used across conditions. Events corrupted by motions (epochs where amplitudes of subpixel rigid registration were larger than 0.5 pixel) were automatically excluded using motion correction information generated in pre-processing procedures.

##### Quantification of spatiotemporal properties

The following statistics were automatically quantified within the GUI: 1. Event frequency (number of spontaneous bands per minute) 2. Event duration (mean duration of spontaneous bands, defined as the half-width of temporal peaks) 3. Inter-peak-interval (IPI, average interval between two neighboring peaks). Note that multiple peaks from the same frames were counted as a single event to exclude zero-length intervals. 4. Normalized bandwidth (defined as the half-width of spatial peaks normalized by the width of the IC) 5. Peak amplitude (average fluorescent intensities at peaks).

##### Global bilateral correlation analysis

Left and right hemispheres of inferior colliculi were manually delineated in the GUI. Mean hemispherical fluorescent traces (averaged over pixels in each hemispheres) were plotted and compared in the GUI (see Supplementary Figure 1C and Figure 1I). Partial correlation between the two average traces was defined as global bilateral correlation and was computed with built-in MATLAB function “partialcorr” controlling the mean fluorescence trace over all pixels outside the inferior colliculi. This analysis was controlled for motions as described before (see Seed-based correlation analysis, this paper).

##### Manual event-labeling

Pre-processed movies were manually assessed in the Manuvent GUI. A tester was asked to label the first and last frames of events by clicking at the best-estimated centers of spontaneous bands up to the human-eye threshold. The tester was trained on a separate set of five movies that were not included for final analysis until the resulting statistics (number of events per minute, and mean event duration) stabilized after labeling the same set of movies 6 times in random order. The GUI allowed users to play movies forward/backward, frame-by-frame, or check/edit labeled events interactively. Movies acquired in different experimental conditions were shuffled and the tester was blinded to any prior information (age, genotype, weight, sex). Summary statistics was based on manually labeled events in both hemispheres of inferior colliculi.

#### Graphic-user-interface For Seed-based Correlation Analysis

A novel Graphic-user-interface (“PickSeedMap” GUI, see Supplementary Figure 4B) was designed for correlation map visualization, representative maps selection, and analyzing regional properties of correlation patterns. Seed-based correlation matrices were generated with the pipeline mentioned above (see Imaging Processing, this paper).

##### Selection of representative seeds

Three representative seeds corresponding to low-/mid-/high-frequency regions were chosen based on typical correlation patterns seen in P3-P7 control animals. Summarized criteria were used across conditions: 1. The first representative seed (corresponding to the low-frequency region) was selected when only a single high-correlation band was visible in each hemisphere (see Figure 1J-K and Movie S3). 2. The next representative seeds were chosen 3-4 seeds down relative to the previous ones where a pair of high-correlation bands were visible in each hemisphere (see Figure 1J-K and Movie S3). 3. Representative seeds were all located in the medial region of left IC. For experiments with large field of view (simultaneous A1/IC imaging), a single representative seed in the mid-/low-frequency region of the left IC was selected for quantifying IC-contralateral-IC, IC-ipsilateral-A1, IC-contralateral-A1 correlations. Another representative seed in the left A1 was selected for quantifying A1-contralateral-A1 correlation (Figure 2H-I).

##### Determine maximum correlation in a region

To determine the maximum correlation value in a region of interest (usually the contralateral hemisphere) with respect to a given seed, a large ROI was defined symmetrically (see Figure 1J-K and supplementary Figure 4A). The pixel within the ROI whose immediate neighborhood hosted the highest mean correlation (using a 5×5 averaging filter) was identified, and this mean correlation value was defined as the maximum correlation in the ROI w.r.t the seed.

##### Compute regional properties of correlation patterns

The Nearby region of the first representative seed (corresponding to the low-frequency region) was converted to a connected component using a threshold of 0.95 (selecting pixels that had >0.95 correlation w.r.t. the seed). The connected component was approximated as an ellipse (see Figure 2D: left panel), and its regional properties were extracted using a built-in function “regionprops” in MATLAB. Eccentricity of this region was quantified as the aspect ratio of the approximated ellipse (a/b). The proportion was calculated as the area of the region normalized by the area of the hemisphere.

#### Determine Auditory Thresholds From Calcium Data

Animals were presented with 6-10 sessions of 80 different frequency-SPL combinations of SAM tones (see Acoustic Stimulation For In vivo Calcium Imaging, this paper). All movies were pre-processed with the same object-oriented pipeline mentioned above (see Image Processing, this paper) leaving out the normalization (ΔF/F0) step. Trials in different sessions but presented with the same frequency-SPL tone were sorted to the same group. Frame-wise averages were computed across all trials within the group. The pre-processing pipeline outputted 80 mean-response matrices/movies (each contained 50 averaged frames). Normalization (ΔF/F0) was conducted after getting the mean-response matrix and F0 was defined as the average fluorescent value of the first 10 frames (before acoustic stimuli were presented) for each pixel. Using the “Manuvent” GUI (see Graphic-user-interface For Wide-field Auditory Data Analysis, this paper), the pixel that had maximum ΔF/F0 values between frame #11-15 (during which the acoustic stimuli were presented) was automatically detected (see Supplementary Figure 10B, left panel: the red dot denoted the max-responding pixel). To obtain a robust response curve based on population activity, a seed-based correlation map was generated with respect to the max-responding pixel using the “PickSeedMap” GUI (see Graphic-user-interface For Seed-based Correlation Analysis, this paper). Populational activity was pooled from pixels within the high-correlation (>0.997) region (see Supplementary Figure 10B, right panel: the region in white). The mean response curve was defined as the average fluorescent trace across all pixels in the high-correlation region (see Supplementary Figure 10B). Animals were considered responding to a tone at certain SPL level if the curve crossed 2% ΔF/F0 threshold during frame #11-20 (see Supplementary Figure 10C). The lowest SPL at which animals responded to the tone was defined as the auditory threshold of the tone. The same criteria were applied across conditions in this study to determine animals’ auditory thresholds. **Generate average response images:** Example response images shown in Figure 5C and supplementary Figure 10C were averaging over frames #11-15 from all sessions during which acoustic stimuli were presented.

#### Graphic-user-interface For Dimensionality Reduction and Unsupervised Clustering

A novel Graphic-user-interface (“GUI_dimReduction”, see Supplementary Figure 3A) was designed for dimensionality reduction, data visualization, and unsupervised clustering of wide-field calcium imaging data based on two commonly adopted techniques: diffusion map (Coifman and Lafon, 2006) and t-SNE (van der Maaten and Hinton, 2008).

##### Dimensionality reduction and data visualization

Pixels in an input movie were projected to and visualized in three-dimensional space by a diffusion map and/or t-SNE based on similarities between their activity patterns (fluorescent traces). For diffusion map analysis, sigma values that determined diffusion speeds on Markov networks was defined as two times the standard deviation of pairwise distances among all pixels (Euclidean distances between pixels in the activity space) in order to adapt with different data landscapes. Key parameters used in t-SNE analysis were consistent across all conditions (see Supplementary Figure 3A. An auxiliary GUI (“CorrespondMaps”) allowed users to correspond projected points in the low-dimensional space back to the original pixels interactively (using built-in brushing function in MATLAB, see Supplementary Figure 3B).

##### Unsupervised clustering

Pixels on line-scans (see Line scan and peak detection, this paper) or movies masked with rectangular ROIs were assessed. K-means unsupervised clustering was performed on the ΔF/F0 traces using MATLAB’s built-in kmeans++ algorithm (Arthur and Vassilvitskii, 2007) with distance metric set as “correlation”. Possible number of clusters (K) ranged from 1-15 (or 1-30) for shuffled data. To generate shuffled data, temporal orders of fluorescent traces of all input pixels were randomized. Best number of clusters was determined with the Davies-Bouldin criterion (the optimal K that minimized the ratio of within-cluster and between-cluster distances) (Davies and Bouldin, 1979). Clustering results could be visualized in the CorrespondMaps GUI. Evaluation were conducted 10 times with randomized initial conditions. The most common number (mode) seen in the 10 repeats was considered the optimal.

## KEY RESOURCES TABLE

| REAGENT or RESOURCE | SOURCE | IDENTIFIER |
| --- | --- | --- |
| **Antibodies** |  |  |
| Anti-Choline Acetyltransferase (ChAT) Antibody (Goat) | Millipore | Cat# AB144P<br>RRID: AB_2079751 |
| Alexa Fluor® 647 AffiniPure Donkey Anti-Goat IgG (H+L) | Jackson ImmunoResearch | Cat# 705-605-147<br>RRID: AB_2340437 |
| **Bacterial and Virus Strains** |  |  |
| rAAV2-hSyn-DIO-hM4D(Gi)-mCherry | Addgene | #44362-AAVrg |
| rAAV2-hSyn-DIO-hM3D(Gq)-mCherry | Addgene | #44361-AAVrg |
| **Chemicals, Peptides, and Recombinant Proteins** |  |  |
| Apamin | Tocris | Cat# 1652<br>CAS# 24345-16-2 |
| alpha-Bungarotoxin | Abcam | Cat# ab120542<br>CAS# 11032-79-4 |
| VECTASHIELD Antifade Mounting Medium with DAPI | Vector Laboratories | Cat# H-1200, RRID: AB_2336790 |
| Clozapine N-oxide | Tocris | Cat# 4936 |
| **Experimental Models: Organisms/Strains** |  |  |
| Mouse: B6.Cg-Snap25tm3.1Hze/J (SNAP25-G6s) | Jackson Laboratory | RRID:IMSR_JAX:025111 |
| Mouse: a9a10 nAChR Knokouts | Morley et al. (2017) | N/A |
| Mouse: B6N.129S6(B6)-Chattm2(cre)Lowl/J (ChAT-Cre) | Jackson Laboratory | RRID:IMSR_JAX:018957 |
| **Oligonucleotides** |  |  |
| **Software and Algorithms** |  |  |
| MATLAB | MathWorks | RRID:SCR_001622 |
| ImageJ | NIH | RRID:SCR_001935 |
| ZEN Digital Imaging for Light Microscopy | Zeiss | RRID:SCR_013672 |
| Behavioral Auditory PHySiology (BAPHY) | Open-source software from the Neural Systems Laboratory, University of Maryland, College Park | https://bitbucket.org/lbhb/baphy/src/master/ |
| BioSigRP | Tucker-Davis Technologies | RRID:SCR_014590 |
| SPIKE2 | Cambridge Electronic Device | RRID:SCR_000903 |
| Object-oriented pre-processing pipeline for wide-field auditory data analysis | This Paper | https://github.com/CrairLab/Yixiang_OOP_pipeline |
| Graphic-user-interface for wide-field auditory data analysis | This Paper | https://github.com/CrairLab/Manuvent |
| Graphic-user-interface for seed-based correlation analysis | This Paper | https://github.com/CrairLab/PickSeedMap |
| Graphic-user-interface for dimensionality reduction and unsupervised clustering | This Paper | https://github.com/CrairLab/GUI_dimReduction |
| Other |  |  |
| Master 8 (multi-channel neurophysiological stimulator) | A.M.P.I. | www.ampi.co.il |
| USG Electret Ultrasound Microphone and UltraSoundGate 116Hb | Avisoft Bioacoustics | part # 40001 |
| Adobe Illustrator | Adobe | RRID:SCR_010279 |
| GraphPad Prism | GraphPad | RRID:SCR_002798 |

## Notes

### Competing Interest Statement

The authors have declared no competing interest.

### Summary of Updates

Adjust pdf format.

